# Benchmarking of SpCas9 variants enables deeper base editor screens of *BRCA1* and *BCL2*

**DOI:** 10.1101/2021.08.18.456848

**Authors:** Annabel K Sangree, Audrey L Griffith, Zsofia M Szegletes, Priyanka Roy, Peter C DeWeirdt, Mudra Hegde, Abby V McGee, Ruth E Hanna, John G Doench

## Abstract

Numerous rationally-designed and directed-evolution variants of SpCas9 have been reported to expand the utility of CRISPR technology. Here, we benchmark PAM preferences, on-target activity, and off-target susceptibility of 11 variants of SpCas9 in cell culture assays with thousands of guides targeting endogenous genes. To enhance the coverage and thus utility of base editing screens, we demonstrate that the SpCas9-NG and SpG variants are compatible with both A>G and C>T base editors, more than tripling the number of guides and assayable residues. We demonstrate the performance of these technologies by screening for loss-of-function mutations in *BRCA1* and Venetoclax-resistant mutations in *BCL2*, identifying both known and new insights into these clinically-relevant genes. We anticipate that the tools and methodologies described here will facilitate the investigation of genetic variants at a finer and deeper resolution for any locus of interest.

## INTRODUCTION

Coupling CRISPR technology to highly parallel methods to write and read DNA^1–3^, as well as viral technologies that enable delivery to nearly any cell type of interest, has enabled pooled genetic screens across diverse models, assays, and fields of study^4^. However, limitations of specificity and activity have inspired researchers to develop variants of the *S. pyogenes* Cas9 (SpCas9) enzyme. Off-target activity may occur at unintended genomic sites, either due to recognition of an alternative protospacer adjacent motif (PAM) site or tolerance of mispairing between the guide and the DNA^5–7^. The frequency of these off-target events is of special concern when developing CRISPR technology for therapeutic applications^8^. As a result, many groups have developed high-fidelity versions of SpCas9. SpCas9-HF1^9^, HypaCas9^10^ and eSpCas9-1.1^11^, developed by rational design, and HiFi Cas9^12^ and evoCas9^13^, identified via randomized screening in bacteria and yeast, respectively, have all been shown to mitigate off-target cutting without substantial loss of on-target activity.

PAM availability also poses another constraint of SpCas9: the canonical NGGN PAM appears approximately every 8 nucleotides, which is sufficient for the knockout of most protein-coding genes, but can be limiting when the location of the perturbation is critical. For CRISPR activation (CRISPRa) and interference (CRISPRi) approaches, the optimal targeting occurs in a relatively narrow window of 50 - 100 nucleotides (nts)^14–16^. The target space is even more limited for homology-directed repair^17^, base editing^18, 19^, and prime editing^20^, for which the maximal activity window is approximately 10 nts. Motivated by this limitation, several groups have created PAM-flexible variants that expand the targeting scope of the already well-characterized SpCas9 enzyme. SpCas9-VQR and SpCas9-VRER, both the result of directed evolution in bacteria, are characterized as recognizing NGA or NGCG PAMs, respectively^21^. SpCas9-NG was rationally engineered to recognize NG PAMs^22^ and xCas9-3.7, identified by phage-assisted continuous evolution (PACE), has been reported to recognize NG, NNG, GAA, GAT and CAA PAMs^23^, although the generalizability of xCas9 has been called into question^24, 25^. More recently, SpG was also developed to recognize NG PAMs, while SpRY has been characterized as essentially PAM-less^26^. Together, these variants have enabled targeting of genomic loci previously inaccessible by wildtype (WT) SpCas9.

Several groups have benchmarked these variants in large scale assays. One study profiled HypaCas9, eSpCas9-1.1 and Cas9-HF1 using a tagmentation-based tag integration site sequencing (TTISS) approach^27^, and found that there is a trade-off between specificity and activity. Notably, this study included only sgRNAs containing 5’ matched guanines, as several groups have reported that these high-fidelity variants do not tolerate a mismatched 5’ guanine^9, 10, 28^, which is often prepended to enhance transcription from the eukaryotic U6 promoter^29^. Zhang and colleagues employed transient transfection and a high throughput sequencing approach to analyze the editing efficiency, specificity and PAM compatibility of high fidelity and PAM-flexible variants at multiple target sites^30^. Additionally, both Legut et al.^24^ and Kim et al.^31^ have assayed xCas9-3.7 and Cas9-NG alongside WT-Cas9 in pooled screens. The former approach used a flow cytometry-based assay in which the authors targeted three cell-surface genes with guides using all possible 3 nucleotide PAMs, the latter used a library on library approach to profile PAMs up to 5 nucleotides. Both found that Cas9-NG is more active at NGH sites than WT-Cas9, but that PAM flexibility comes at the cost of reduced efficacy^24^.

Previously, we and others have demonstrated the utility of screens using base editor technology to introduce variants at their endogenous loci^32, 33^, identifying loss-of-function mutations in clinically-relevant genes, as well as variants that modify the ability of small molecules to interact with their targets. We expand upon existing work with the goal of assaying any new SpCas9 variant, either high-fidelity or PAM-flexible. After identifying two useful PAM variants that perform well, Cas9-NG and SpG, we develop these enzymes for base editing applications. We build off our previous work with BE3.9 for C>T editing^33^ and assay the performance of a recently described A>G editor (ABE8e)^34^ for use in pooled screens. We show that such screens can uncover loss-of-function mutations in *BRCA1* by dropout screens, and as we demonstrate with Venetoclax and *BCL2*, are particularly powerful for mapping drug - target interactions by resistance screening.

## RESULTS

A new Cas protein, whether an engineered variant of an already-described Cas protein or isolated from a novel source, can be characterized by its on-target activity - how efficiently it targets its intended sequence - and its propensity for off-target activity - cleavage at unintended sites. Affecting both of these properties is the PAM sequence preference of each Cas protein, that is, which PAMs are consistently active at on-target sites, and which have lower efficiencies. For this latter group of PAMs, the exact point at which they become detrimental, meaning their potential for off-target activity surpasses their on-target utility, remains to be well-characterized. To better quantify these performance metrics among SpCas9 proteins, we began by designing a PAM-mapping library that reports on both the PAM preferences and on-target efficacy of any SpCas9 variant, based on its ability to distinguish between thousands of essential^35^ and nonessential genes^36^. The library contains 70 - 100 sgRNAs per four nucleotide PAM, including all 256 possible PAMs (the canonical SpCas9 PAM is thus NGGN). Further, each of these 70 - 100 sgRNA sets includes three different sgRNA 5’-types which differ in length and/or the presence of a matched 5’ guanine: G19, a 20mer with a matched guanine; G20, a 21mer with a matched prepended guanine; and g20, a 21mer with a mismatched prepended guanine (**Fig 1a**). We used a consistent lentiviral vector architecture, in which the expression of SpCas9 is driven by the EF1a promoter, and introduced point mutations in Cas9 to create nine different variants as well as unmodified (WT) Cas9. We first established A375 (melanoma) cells stably expressing these 10 constructs and selected with blasticidin for 14 days. We next introduced the PAM-mapping library, which has 18,768 total guides, into each of the 10 pre-established lines in duplicate, selected with puromycin for 5 - 7 days, and maintained each population with at least 500x coverage for an additional two weeks (**Fig 1b**). At the end of the screen, we collected cells, isolated genomic DNA, retrieved the library by PCR, and performed Illumina sequencing to determine the abundance of each guide.

**Figure 1.**
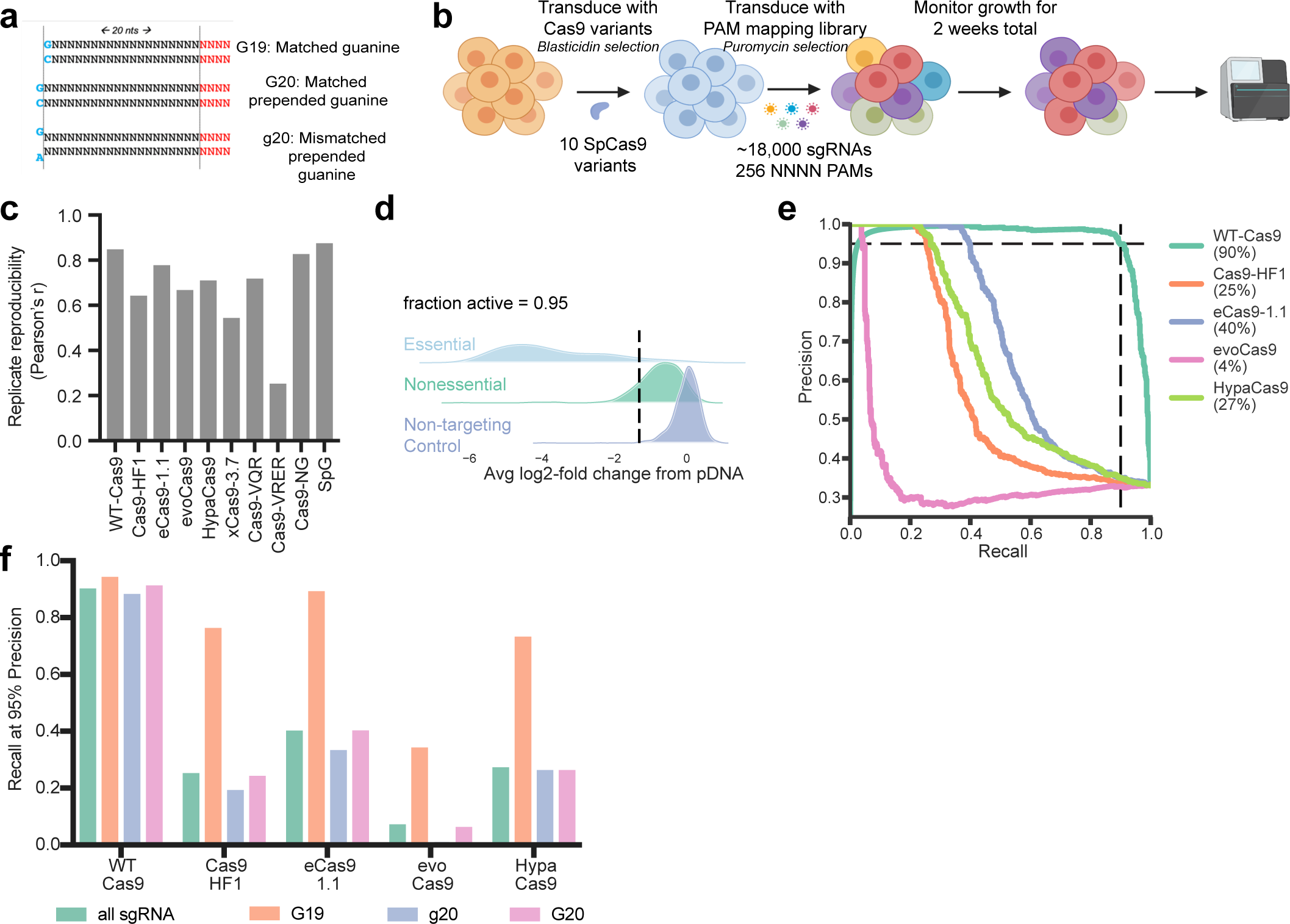
Establishment of a benchmarking assay for Cas9 activity. a) Schematic of PAM-mapping screens. b) Schematic of the three 5’-sgRNA types screened. Location of the PAM sequence is indicated in red. c) Replicate correlation (Pearson’s r), calculated from n=2 experimental replicates for each variant screened. d) Example fraction active calculation for WT-Cas9 at NGGN PAMs. e) Precision-recall curves for WT-Cas9 and high fidelity variants profiled with the PAM-mapping library. Guides of all 5’-types are included in this calculation. Dashed lines designate the recall at 95% precision for WT-Cas9. f) Recall values at 95% precision for WT-Cas9 and high fidelity variants profiled with the PAM-mapping library (NGGN PAMs only), discretized by 5’-sgRNA type.

To interpret the results, we first calculated the log2-fold-change (LFC) compared to the initial library abundance, as determined by sequencing the plasmid DNA (**Supplementary Data 1**). Replicates had a wide range of Pearson correlations (0.25 to 0.87); generally, lower-replicating variants had few active guides in this assay (**Fig 1c**). We quantified the fraction of guides targeting essential genes that were more depleted than the 5th percentile of guides targeting non-essential genes and non-targeting controls for each of the 256 PAMs assayed with each variant. For WT-Cas9, this revealed the expected preference for an NGGN PAM, for which 95.3% of guides were active by this metric (**Fig 1d**), with low but detectable activity at NAGN (18.6%), and minimal activity above background at NGAN (6.1%) and NCGN (4.7%) PAMs (**Supplementary Figure 1a)**. We examined the data via an alternative metric, calculating average recall at 95% precision for each variant, designating guides targeting essential genes as true positives and those targeting nonessential genes as false positives. We found that these metrics produce concordant results (Pearson’s r = 0.98 for WT-Cas9) (**Supplementary Figure 1b**), with an average recall of 90.3% for an NGGN PAM, 18.7% at NAGN, 5.2% at NGAN, and 4.3% at NCGN PAMs.

### High fidelity variants

#### On-target efficacy

As expected, the high fidelity variants were only active at NGGN PAMs (**Supplementary Figure 1a**), so we included only these PAMs in downstream analyses. We calculated precision-recall curves as above, including guides of all 5’-types (G19, G20 and g20). At 95% precision, WT-Cas9 performed best (90% recall), followed by eCas9-1.1 (40%), HypaCas9 (27%), Cas9-HF1 (25%); and finally evoCas9 (4%) (**Fig 1e**). When we discretized the data by 5’-type, we observed a pronounced preference for G19 guides with all high-fidelity variants, but not with WT-Cas9 (**Fig 1f**). Considering only this subset of G19 guides, WT-Cas9 again had the best recall (94%), followed by eSpCas9-1.1 (90%), Cas9-HF1 (76%), HypaCas9 (74%), and evoCas9, (35%), a ranking consistent with a recent study using reporter construct screens to categorize these variants^25^. That an extra 5’G, whether paired or not, greatly diminishes activity with these variants is likewise consistent with prior reports^27–29^. Importantly, this guide composition constraint reduces the number of potential sgRNAs 4-fold. We did not observe a substantial difference between activity with G20 vs. g20 sgRNAs so we summarized PAM activity into active (guides with fraction active >0.7) and intermediate (fraction active 0.3 - 0.7) bins for 21mers (G20/g20) and 20mers (G19) for these variants at all PAMs (**Supplementary Figure 1c**).

#### Off-target activity

We next sought to compare the off-target tolerance of select high fidelity variants to that of WT-Cas9. To systematically assess off-targets based on mismatches in the sgRNA sequence, we collated a set of 21 sgRNAs that were active with every high fidelity variant in the PAM-mapping assay (all G19 guides), and included all possible single (n = 1,197) and double mismatches (n = 32,319) in the sgRNA sequence, as well as 1,000 non-targeting controls, resulting in a library of 34,537 guides (**Fig 2a**). We performed screens in duplicate in A375 cells stably expressing three different variants: WT-Cas9; eCas9-1.1, as this was the best-performing variant in the PAM-mapping library, and HiFi Cas9, another recently described variant we had not previously assessed^12^. Sequencing the sgRNAs after three weeks of growth, we found that replicates were well correlated (Pearson’s r = 0.90 - 0.93, **Supplementary Data 2**). We determined the LFC of perfect match, single mismatch, double mismatch, or non-targeting control guides and plotted the distribution of guides by type (**Fig 2b**). To quantitate off-target activity, we calculated an ROC-AUC measuring the separation between perfectly matched guides (positive controls) and either single or double mismatched guides (negative controls) (**Fig 2c**).

**Figure 2.**
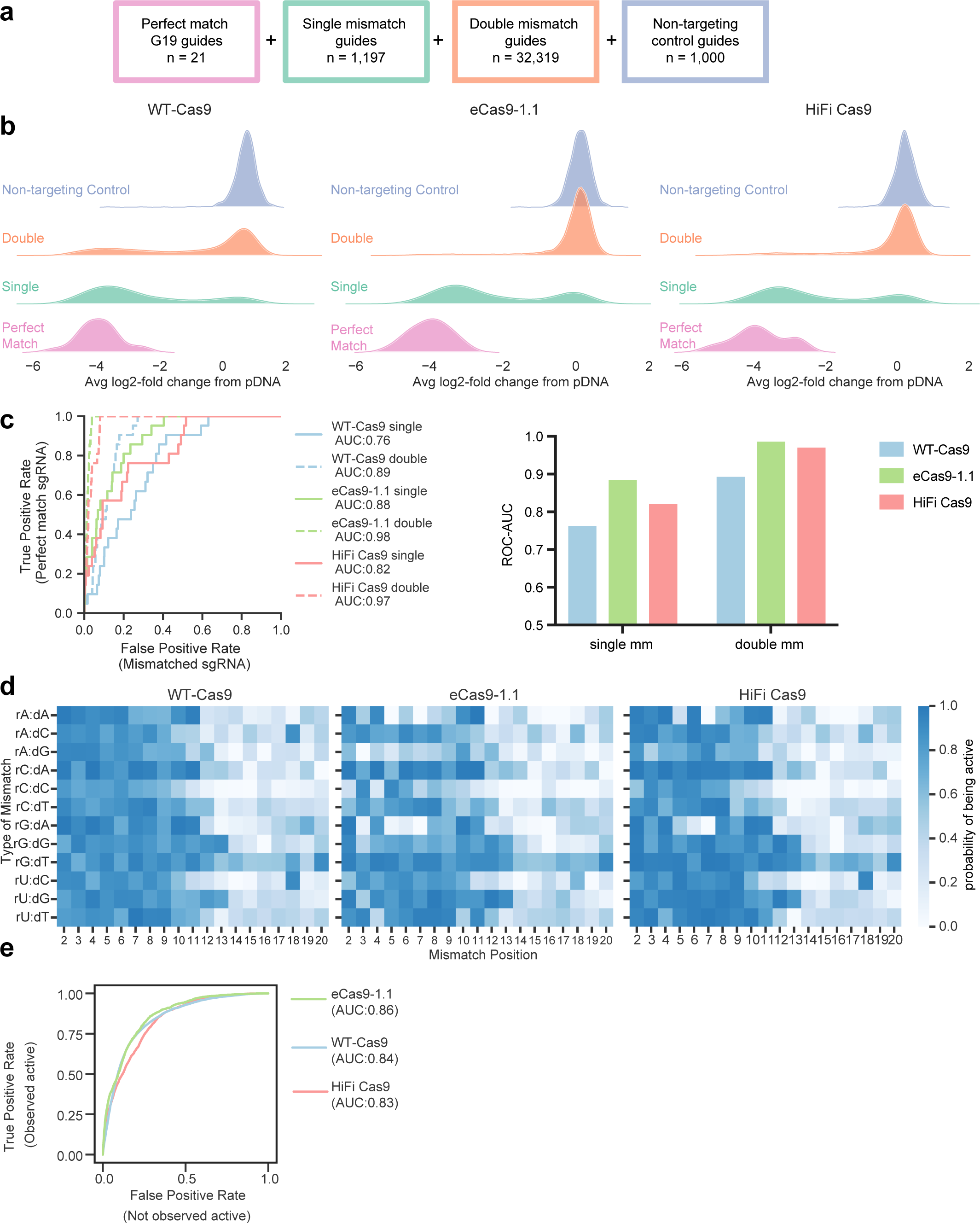
Off-target profiles of high fidelity variants. a) Schematic depicting off-target library construction and guide selection. b) Ridge plots showing activity of guides in the library with zero, one or two mismatches. c) ROC plots for each enzyme screened with single (solid lines) and double mismatched sgRNAs (dashed lines). AUC is reported in the graph legend. Data are also summarized in a barplot. d) CFD matrices for each enzyme, numbered such that 2 is the second nucleotide in the guide. Note that mismatches start at position 2, because the first position of the guide is always fixed as a G. e) ROC plot depicting ability to predict activity at double mismatches using single mismatch data. True positives are guides that were observed to be active, false positives are guides that were were not active in the screen.

We then calculated the probability of being active for each mismatch type and position to generate a cutting frequency determination (CFD) matrix for each variant, as done previously with SpCas9 and AsCas12a^6, 37^ (**Fig 2d**, **Supplementary Table 2**). Here, we used a logistic regression model to transform the LFC values to a probability of being active, defining perfect match sgRNAs as positive controls and non-targeting sgRNAs as negative controls. We had previously generated a CFD matrix for WT-Cas9 using guides mismatched to the gene CD33^6^, and these new results were moderately consistent (Pearson’s r = 0.61, **Supplementary Figure 2a**). Note that there were several experimental differences between these two assays: the present study used only G19 sgRNAs while in 2016 we did not impose a 5’-type requirement; the present study assayed 14 essential genes, while the latter focused on a single gene; and the readouts were different, viability versus flow cytometry, respectively. Here, we observed a higher tolerance for mismatches at the PAM distal end of the guide with all three enzymes, as well as for rG:dT mismatches (**Fig 2d**), two trends observed previously with other techniques to examine off-target activity of SpCas9 and which have also been seen with Cas12a enzymes^5, 6, 37^. We also find that both high fidelity variants show greater discrimination for rG:dA and rA:dA mismatches than WT-Cas9 (**Fig 2d**, **Supplementary Figure 2b**), whereas other mismatches, such as rC:dA and rU:dG, are still substrates for cleavage by these enhanced specificity variants (**Fig 2d**, **Supplementary Figure 2b**).

Using the product of the activities of each individual mismatch in the CFD matrix, we predicted the activity of double-mismatch guides, an approach that has been validated by others to identify problematic off-target sites with more than one mismatch for SpCas9^37–39^. To evaluate our predictions, we drew ROC curves designating positive controls as guides that were observed to be active, and negative controls that were observed to be inactive (**Fig 2e**). We saw good discrimination between the two sets (AUC = 0.83-0.86) for all three enzymes, suggesting that this approach can help identify problematic, multi-mismatch, off-target sites when designing guides.

### PAM-flexible variants

#### On-target activity

Returning to the screens with the PAM-mapping library, we analyzed the activity of SpCas9 variants engineered to recognize alternative PAMs (**Supplementary Figure 1a**). Cas9-VQR was initially characterized as recognizing NGAN PAMs, with a preference for NGAG > NGAA = NGAT > NGAC^21^. Our results are consistent with this initial characterization; we observe excellent activity with all 4 NGAG PAMs (fraction active > 0.9), and diminished activity at the remainder of the NGAN PAMs (0.11 - 0.67) (**Fig 3a, b**, **Supplementary Figure 1a**). The Cas9-VRER variant, characterized to target NGCG PAMs, showed intermediate activity in this assay with GGCG (0.36) and poor activity with HGCG (0.19 - 0.28).

**Figure 3.**
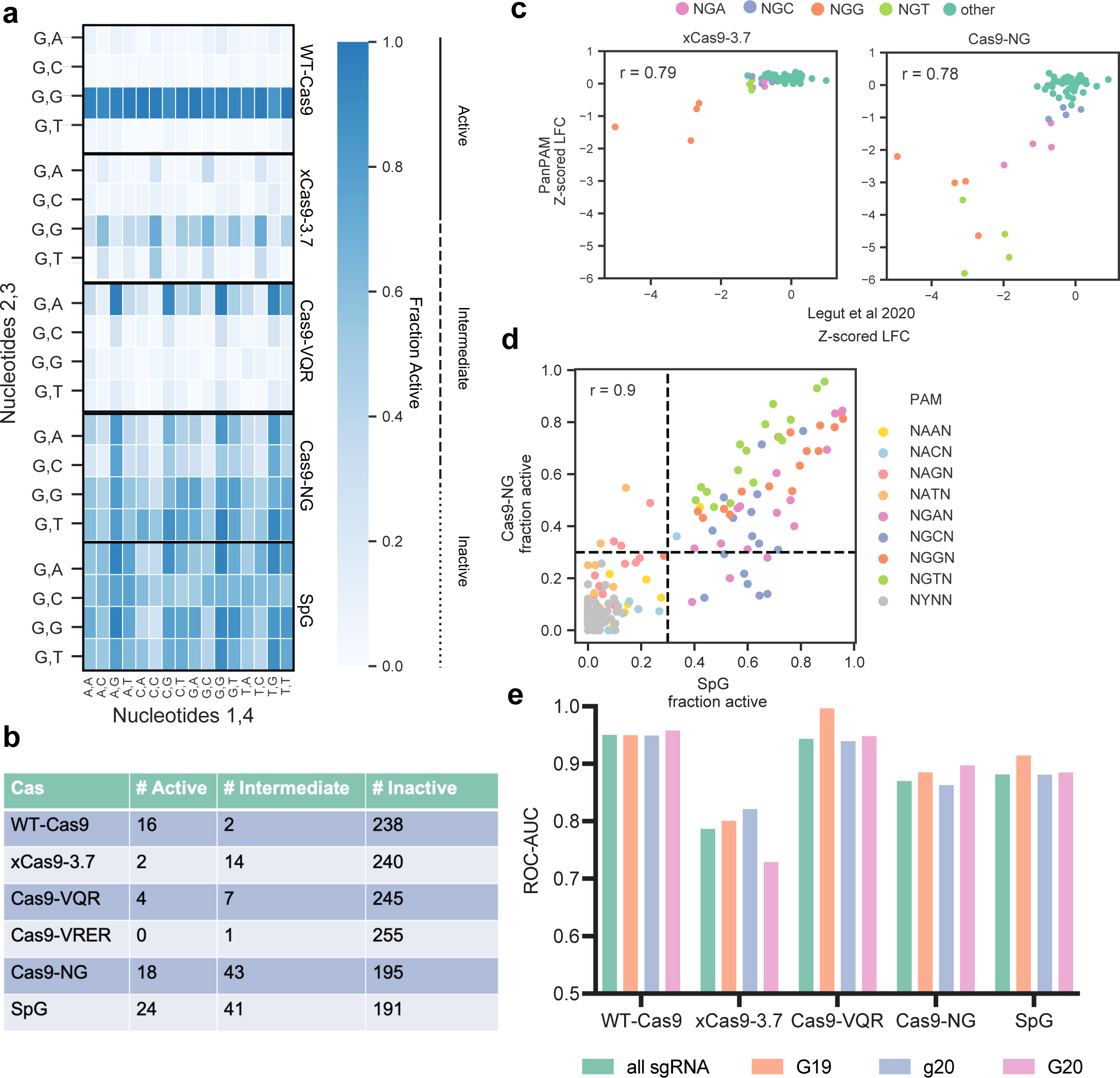
Benchmarking PAM-flexible variants. a) Heatmap of fraction active values at all NGNN PAMs. Nucleotides 1 and 4 are along the x-axis, nucleotides 2 and 3 along the y-axis. b) Table of fraction active values for each PAM-flexible variant binned by activity bin. c) Comparison of xCas9-3.7 (left) and Cas9-NG (right) to Legut et al. 2020. Points are colored by PAM. PAM-mapping z-scored LFC values on the y-axis refer to data in the present study. d) Comparison of Cas9-NG and SpG fraction active values. Points are colored by PAM. Dashed lines at 0.3 indicate the cutoff for intermediate PAMs. e) ROC-AUC values by 5’-type for each PAM-flexible variant. True positives are guides targeting essential genes, false positives are guides targeting nonessential genes. Only active PAMs are considered in this analysis.

For xCas9-3.7, only two PAMs showed high activity, both of which were canonical NGGN sites (CGGC and TGGC), with 9 additional NGGN PAMs showing intermediate activity (fraction active 0.35 - 0.66), and the remaining 5 NGGN showing low activity (0.16 - 0.29). We identified 5 additional PAMs with intermediate activity (4 NGTN, 1 NGAN) for a total of 14 intermediate PAMs (**Fig 3a,b**). Legut et al. recently characterized xCas9-3.7 at all possible 64 NNN PAMs using a flow cytometry-based assay^24^. We z-scored sgRNAs targeting the coding regions of CD45 and CD55 used in their assay, and observed concordance with essential sgRNAs from the present study at all PAM sites (Pearson’s r = 0.79), with the majority of sgRNAs centered around 0, and activity only at NGG PAMs (**Fig 3c**). Further, Kim et al.^31^ performed a similar PAM classification study utilizing a high-throughput reporter assay to measure indel frequencies at 4 and 5 nucleotide PAMs. We compared our fraction active metric against their indel frequency using WT-Cas9, and observed good concordance (Pearson’s r = 0.95, **Supplementary Figure 3a)**. For xCas9-3.7, we observed a similar trend (Pearson’s r = 0.90), with the vast majority of PAMs centered around 0, and the strongest activity at NGGN sites, with modest activity at some NGHN sites (**Supplementary Figure 3b**).

In contrast to the poor activity of xCas9-3.7, we identified 18 active PAMs with Cas9-NG^22^, including high activity at NGTG and NGAG PAMs, but diminished activity at NGAC and NGCC PAMs (**Fig 3a**), consistent with prior results^22, 24, 31^. We observed intermediate activity at 43 additional PAMs (**Fig 3b**). Using the same comparison metric as above, we observed a similarly strong correlation between Cas9-NG in our assay and the results from Legut et al. (Pearson’s r = 0.78) (**Fig 3c**). We also observed concordance between our fraction active metric and indel frequency measurements from Kim et al. with Cas9-NG (Pearson’s r = 0.84, **Supplementary Figure 3c**). Interestingly, xCas9-3.7 and Cas9-NG, which were both described as recognizing NG PAMs, show little correlation at these PAMs in our assay (Pearson’s r = 0.24, **Supplementary Figure 3d)**, which differs from the relationship observed by Legut et al. (Pearson’s r = 0.72, **Supplementary Figure 3e**).

Finally, we identified 24 active PAMs with the SpG variant, all of which were NGNN, consistent with the initial characterization^26^ (**Fig 3a**). An additional 41 PAMs showed intermediate activity, 39 of which were NGNN and the remaining 2 NANN (**Fig 3b**). We next compared Cas9-NG with SpG, as these enzymes have similar expanded PAM profiles. We observed good concordance on the guide level (Pearson’s r = 0.79 with all sgRNAs, r = 0.82 when filtered for NG PAMs) (**Supplementary Figure 3f, g**), and on the PAM level (Pearson’s r = 0.9) (**Fig 3d**). We found that some PAMs were more active with SpG than with Cas9-NG, with a few exceptions; Cas9-NG had more activity with NANN PAMs than SpG (**Fig 3d**, **Supplementary Figure 1a**).

To understand if these variants had any 5’ sgRNA-type requirement, we generated ROC-AUC curves for each enzyme for each 5’-type, filtering on active PAMs for each enzyme, and designating guides targeting essential genes as true positives, and guides targeting nonessential genes as true negatives (**Fig 3e**). Consistent with Kim and colleagues’ findings^31^, we found that none of these PAM-flexible variants demonstrate a marked preference. This broad sgRNA compatibility is particularly attractive for modalities limited by targetable sites, such as base editing.

#### Off-target profiles of Cas9-NG and SpG

To characterize the tolerance of SpCas9-NG and SpG for guide-target mismatches, we first identified active, perfect-match sgRNAs from the original PAM-mapping screens and selected a random subset containing 300 of these sgRNAs, maintaining the balance across different PAM sites. We then generated all possible single mismatches and a random subset of double mismatches, resulting in a library with 300 perfect match guides, 17,775 single mismatches, 60,000 double mismatches and 1,000 non targeting controls (total size = 79,075) (**Fig 4a**). In this library, all three 5’-sgRNA types were included. We screened this library in duplicate in A375 cells stably expressing Cas9-NG or SpG.

**Figure 4.**
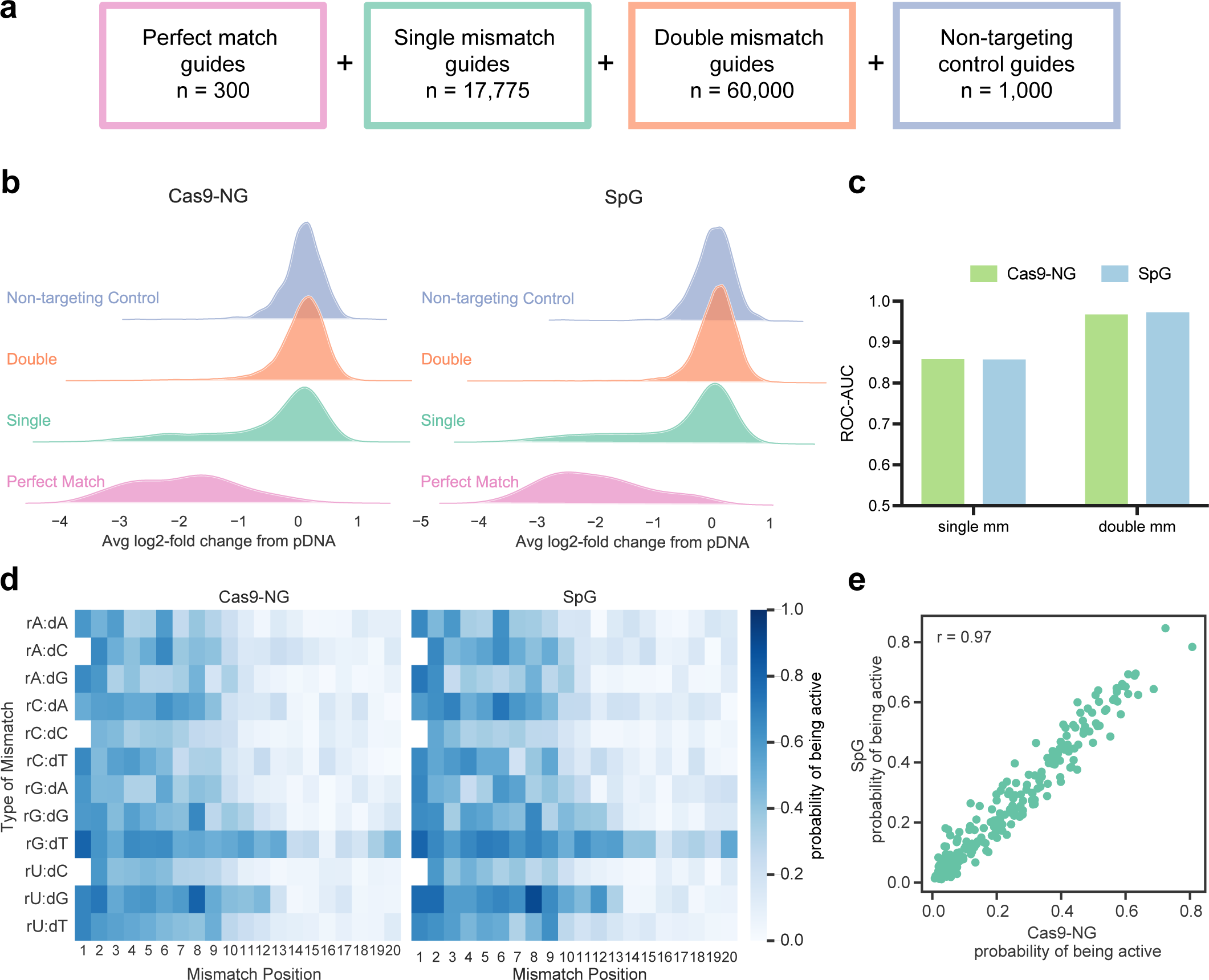
Off-target profiles of Cas9-NG and SpG. a) Schematic depicting off-target library construction and guide selection. b) Ridge plots showing activity of filtered guides with zero, one or two mismatches. c) ROC-AUC values at single and double mismatches for Cas9-NG and SpG. d) CFD matrix for Cas9-NG and SpG. Note that there are no wildtype g20 and G20 guides with a G in the first position, so the rN:dC squares are blank. e) Scatter plot showing probability of being active for each single mismatch position/type for Cas9-NG and SpG (n = 237). Pearson correlation is noted in the top left.

After sequencing the sgRNAs, we calculated LFC relative to pDNA, and found that replicates were well-correlated (Pearson’s r = 0.81 Cas9-NG; r = 0.76 SpG; r = 0.72 Cas9-NG vs SpG, **Supplementary Figure 4a, Supplementary Data 3**). We examined the LFC of perfect match, single mismatch, double mismatch, or non-targeting control guides, considering every guide included in the library (**Supplementary Figure 4b**). From there, we selected 149 of the original 300 perfect match guides that showed the highest activity for subsequent analyses (**Fig 4b**). We applied the same framework of assessing off-target activity as before by calculating the ROC-AUC, comparing single and double mismatched guides to perfect matches (**Fig 4c**). We observed good separation between perfect matches and single mismatches (Cas9-NG AUC = 0.86; SpG = 0.85) and excellent differentiation between perfect matches and double mismatches (Cas9-NG = 0.97; SpG = 0.97).

We then calculated the probability of being active for each enzyme with each mismatch type and position using all 5’-types (G19, G20, g20) to generate a cutting frequency determination (CFD) matrix (**Fig 4d**). We also calculated CFD matrices separated by 5’-type and compared the probabilities of being active by mismatch type and position across guide types within each enzyme (**Supplementary Figure 4c, d**). For both enzymes we found that g20 guides are the least prone to off-target cutting, followed by G20 and finally G19 guides. Finally, we compared the probability of being active for Cas9-NG and SpG by type of mismatch using all 5’-types and observed excellent concordance (Pearson’s r = 0.97) (**Fig 4e**). Cas9-NG and SpG have 7 and 6 mutations in total, respectively, 4 of which are at the same residues, and one of which is the identical substitution (T1337R, **Supplementary Figure 4e**). Given these similarities, it is unsurprising that these variants behave so similarly.

### Base editing with PAM-flexible variants

A major appeal of these PAM-flexible variants is their potential for use in base editor screens, where the location of the perturbation is crucial for introducing the precise desired edit. Likewise, while we have previously demonstrated the utility of C>T base editors (CBEs) in pooled screens, additional base editors capable of altering other nucleotides, such as A>G base editors (ABEs), would further expand the utility of such screens.

We benchmarked 3 different versions of ABEs: ABE7.10^19^ and the more recently described ABE8e^34^ and ABE8.17^40^ in a small-scale assay using a reporter construct containing two sgRNAs targeting EGFP and EGFP itself, delivered via lentivirus to MELJUSO cells (**Supplementary Figure 5a)**. After Sanger sequencing the target site, we quantitated the nucleotide percentage at each editable A (A5 and A8 with EGFP sg1 and A4 and A9 with EGFP sg2) using EditR^41^. We observed the most efficient editing with ABE8e constructs with both guides at all four editable adenines (**Supplementary Figure 5b**), so we selected this ABE for further study. We next generated a Cas9-NG version of ABE8e and tested it in the same assay. Editing levels were lower compared to nickase WT-Cas9, but still achieved as high as 56% editing (**Supplementary Figure 5b**).

#### BRCA1

To understand the current applications of base editing screens, using *BRCA1* as an example target, we calculated the number of residues in which one could introduce a missense or nonsense mutation (see Methods). WT-Cas9 is able to target 455 unique residues in the longest isoform of *BRCA1*, or 24.2% of the protein (220 residues are targetable with CBE and 305 with ABE) (**Fig 5a**). With Cas9-NG, considering PAMs characterized above as active or intermediate, the number of targetable residues increases to 771 with CBE and 989 with ABE, for a total of 1,342 unique residues (71.2% of the protein). Likewise, with SpG, 75.3% of the protein can be modified with at least one mutation.

**Figure 5.**
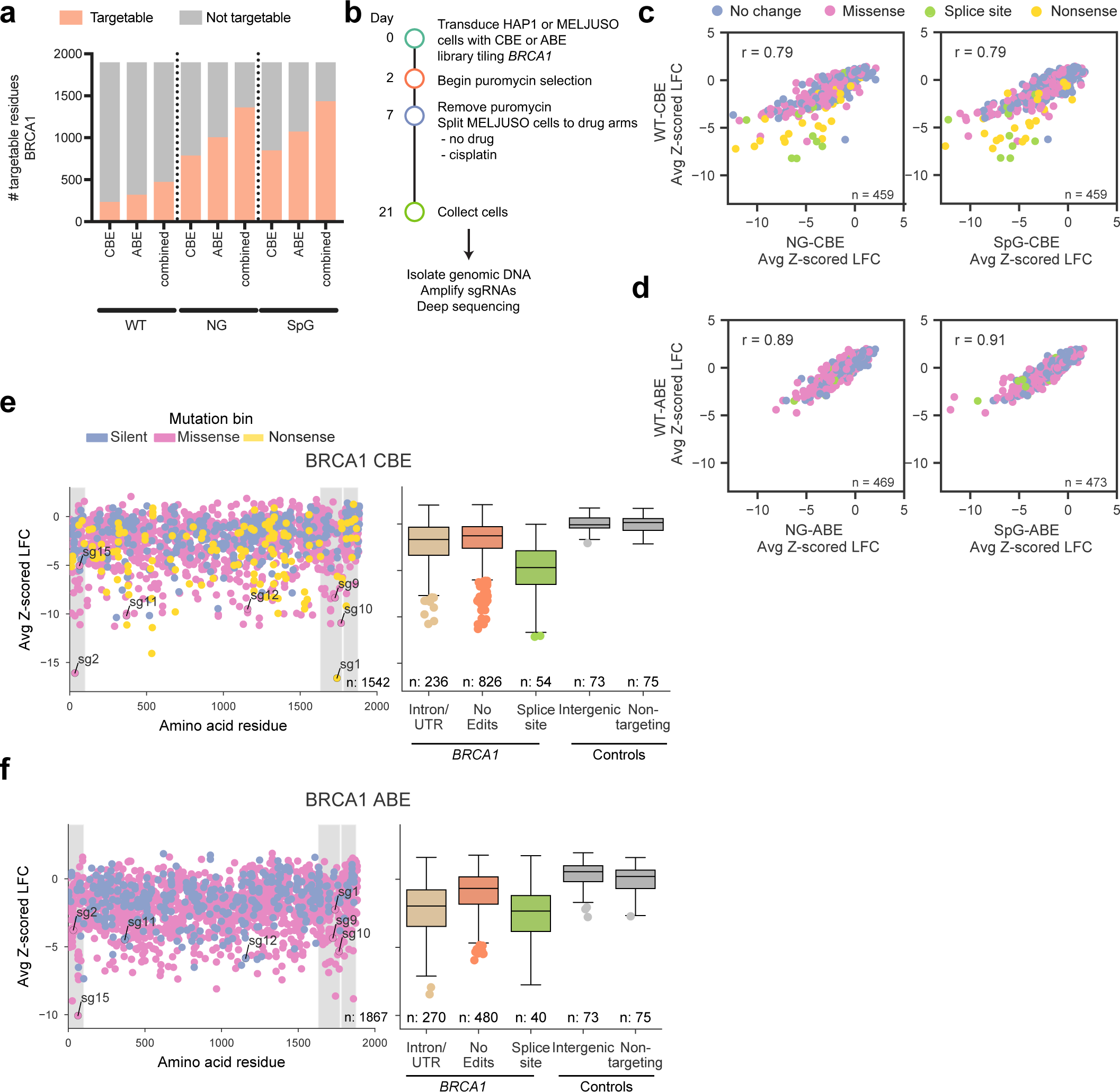
Tiling *BRCA1* with variant base editors. a) Number of targetable residues in BRCA1 using the base editors paired with the library described in this study. b) Timeline by which tiling screens were conducted. c) Comparison of NG and SpG-CBEs to WT-CBE with shared guides predicted to introduce no change (silent or no edits), splice site, nonsense, or missense mutations with an NGGN PAM. Pearson’s r is reported for each comparison. d) Same as (c) but for ABEs. e-f) Average performance of sgRNAs (averaged Cas9-NG and SpG screens) targeting *BRCA1*, colored according to the predicted mutation bin, for CBE and ABE screens. The first grey shaded region spans the RING domain, and the following two indicate the BRCT repeats. Boxes show the quartiles; whiskers show 1.5 times the interquartile range.

Using BE3.9max and ABE8e as the starting vectors (**Supplementary Figure 5c**), we designed two base editor versions of each PAM-flexible variant, which we refer to as NG-CBE, SpG-CBE, NG-ABE, and SpG-ABE. To test these four base editor - Cas variant pairings in a screening context, we designed a library tiling across the DNA-damage repair gene *BRCA1,* containing all possible guides targeting the gene, irrespective of PAM (n = 11,524), and included 30 guides targeting the splice sites of essential genes^35^, 75 intergenic sites, and 75 non-targeting guides. We screened these 4 pairings in 2 cell lines, HAP1 and MELJUSO, in duplicate or triplicate at high coverage (>2,000 cells per sgRNA) for a total of 21 days (**Fig 5b**). Since *BRCA1* is essential in HAP1 cells^42^, we conducted a negative selection (dropout) assay in this cell line, and we treated MELJUSO cells with a low dose (1 µM) of cisplatin^33^ to enhance selective pressure for *BRCA1* loss-of-function alleles. Previously, we had screened *BRCA1* with a tiling library containing only NGG PAMs using a WT-CBE^33^; we re-screened this library with WT-ABE as well.

After calculating LFC values relative to the plasmid DNA, we found that replicates were well-correlated within and between cell lines (Pearson’s r ranging from 0.77-0.99 in CBE screens, **Supplementary Data 5**; 0.77-0.95 in ABE, **Supplementary Data 6**), and thus we averaged the data across the two cell lines. First, we examined the distribution of positive (essential splice sites) and negative (non-targeting and intergenic) controls and found that in all conditions, negative controls were centered around 0, while the positive controls were depleted (**Supplementary Figure 5d**), confirming that there was base editing activity in each screen. To understand our ability to assay *BRCA1* itself with these base editors, we examined the separation of guides predicted to introduce nonsense or splice mutations in *BRCA1* (positive controls) compared to guides predicted to introduce silent or no edits in *BRCA1* (negative controls) and calculated an area under the curve (AUC) for each base editor and cell line (**Supplementary Figure 5e, f**). Across all conditions, we observed the best performance with the WT base editors and slightly higher performance in the near-haploid HAP1 cell line, consistent with our original benchmarking of the base editing technology^33^. In every condition, we observed a clear separation between positive and negative controls, confirming that we were able to assay the *BRCA1* gene effectively. Next, we compared the NG and SpG base editors to WT, using guides with NGGN PAMs targeting the coding sequence of *BRCA1*, and observed good concordance (**Fig 5c, d**), although some guides showed markedly less activity with WT in the CBE arms. We speculate that this relates to the decreased activity seen overall with NG and SpG compared to WT at some NGGN PAMs, which may be especially meaningful for CBE, as continued localization of the UGI domain is necessary for proper base editing. Finally, guides in the library behaved largely similarly when paired with ABE or CBE (Pearson’s r = 0.73 SpG; r = 0.71 NG, **Supplementary Figure 5g**) however there were clear outliers.

We next plotted the average depletion of guides introducing coding changes screened with CBE and ABE along the length of the BRCA1 protein (**Fig 5e, f**). With both technologies, we observed strong depletion of guides targeting the RING and BRCT domains, consistent with our previous findings^33^ and the known clinical importance of these regions. However, it is inappropriate to draw conclusions about specific casual mutations from the behavior of sgRNAs solely from the results of a primary screen, as sgRNAs might deplete due to out-of-window editing, C>R editing, or indels^33^. Further, especially in a negative selection screen, guides can erroneously score due to off-target effects. In order to make conclusions about these hits, we selected several for a focused follow-up.

#### BRCA1 Validation

We selected 18 guides (sg1-18) for focused validation experiments based on the magnitude of z-scores in the primary screen, including guides that depleted despite using an inactive PAM. We cloned individual sgRNAs into either ABE or CBE vectors, transduced cells, and collected samples after one (early time point), two and three weeks post transduction. We then PCR amplified the edited locus using custom primers, and Illumina sequenced the edited genomic loci to identify the causal mutations (**Supplementary Figure 6a**).

Examining the editing efficiency of samples collected at the early time point, we observed a wide range of editing (0.04 - 60.1%, C>T; 0.2-59.1% A>G). In all cases with <1% editing, the sgRNA utilized an inactive PAM, thus confirming that the depletion observed in the primary screen was due to an off-target effect. Of samples with an intermediate or active PAM, there was an average of 43.3% C>T editing and 37.4% A>G editing in the predicted edit window of 4-8 nucleotides, with lower but detectable levels outside of the window (**Supplementary Figure 6b**), consistent with the previously-described properties of these base editors^33, 40, 43^. Next, we examined the reproducibility of percent change with WT alleles, comparing the percentage of reads in the late versus early samples across replicates (**Supplementary Figure 6c**). We found that >10% enrichment of the WT allele was reproducible across replicates, and thus we considered a guide to validate if the WT allele enriched >10% from early to late samples, indicating that edited alleles were depleted. By this criterion, 4 of the 7 guides with an active or intermediate PAM that depleted in the primary screen with CBE validated, and 2 of 5 guides validated with ABE.

For a number of guides we conducted validation studies with both CBE and ABE. When screened with CBE, sg2 generates a P34F mutant, which depletes from 48.0% abundance on day 8 to 11.2% on day 21 (**Fig 6a, b**). Residue 34 makes up part of the RING domain, which forms a heterodimer with the RING domain of BARD1 and is necessary for E3 ubiquitin ligase activity^44, 45^. Although the P34F mutation has not been documented in NCBI’s ClinVar Database, several lines of evidence suggest that this residue plays a critical role. First, it lies in the central RING motif of BRCA1, which is composed of residues 23-76^45^. Additionally, of the 6 possible missense mutations introduced by Findlay et al. via Saturation Genome Editing (SGE), 5 scored as LOF, and 1 as intermediate (P34F requires mutating 2 nucleotides, so was not included in that dataset)^42^. To gain structural insight into the LOF phenotype associated with this mutant, we visualized these residues on the crystal structure of the RING domains of BRCA1 and BARD1 (PDB IJM7). Zn^2+^ atoms stabilize the structure within the RING finger and are maintained by two binding loops, Site I and Site II^45^. P34 falls between Sites I and II on BRCA1, and if mutated to F34, would come into close proximity to C66 on BARD1, which is one of the residues comprising Site II on BARD1 (**Fig 6c**). While further experimentation is required to understand the exact mechanism for this LOF phenotype, it seems likely that the phenylalanine substitution at position 34 disrupts the Zn^2+^ coordination, thereby destabilizing the interaction between BRCA1 and BARD1.

**Figure 6.**
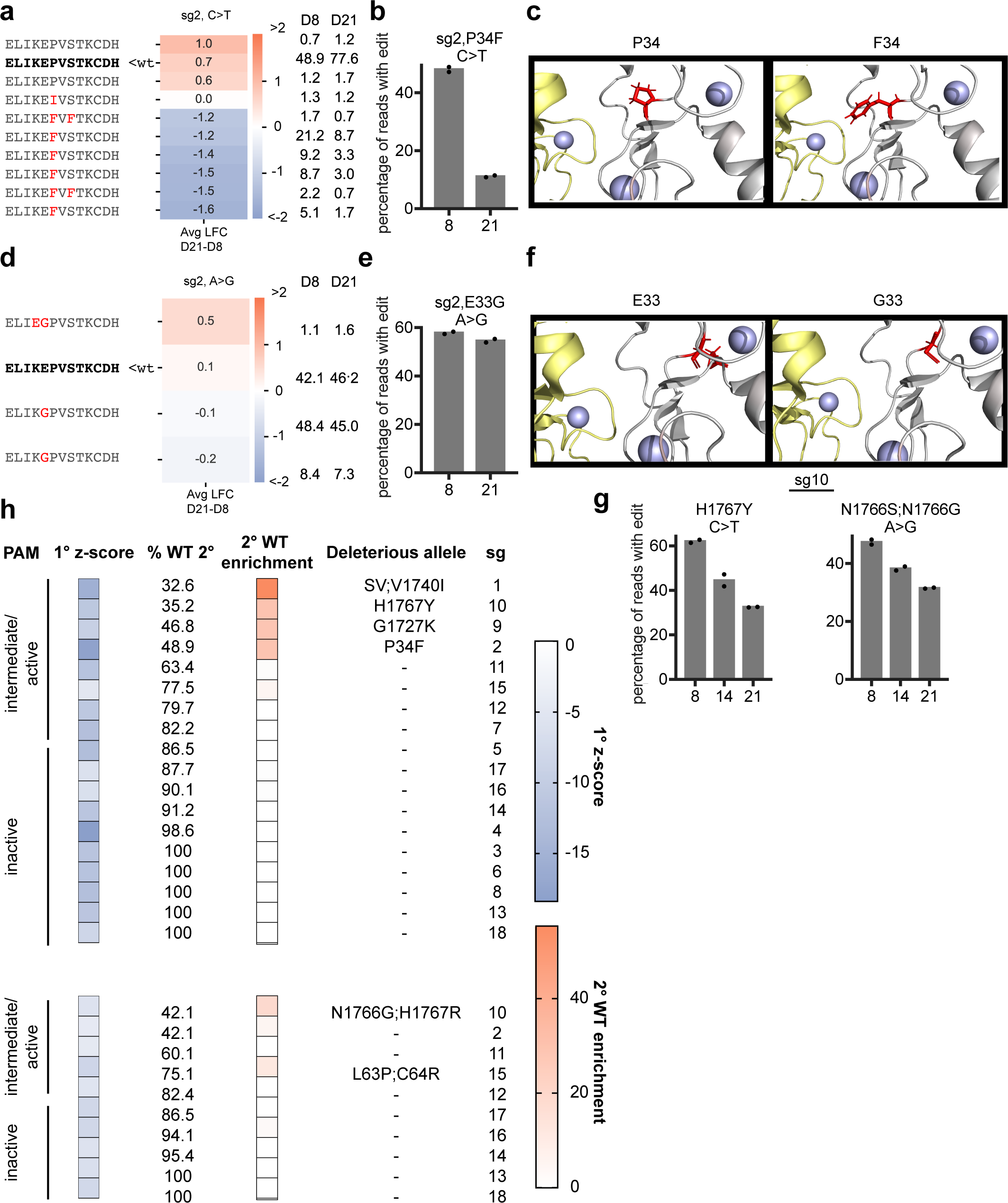
Validation of *BRCA1* hits identified by Cas9-NG. a) Translated sequence around the sgRNA for any allele with at least 1% abundance in any condition. The WT sequence is bolded in black, unchanged amino acids are in grey, and substitutions are highlighted in red. Avg LFC from day 21 - day 8 is indicated on the heatmap and relative percent abundance of each allele is indicated to the right (normalized after filtering for alleles with <1% abundance at both timepoints). b) Percentage of all sequencing reads containing the indicated mutation at each timepoint. Dots indicate n=2 biological replicates. c) View of the RING domain (PDB IJM7) of BRCA1 (grey) bound to the RING domain of BARD1 (yellow), with Zn2+ atoms in purple. The left panel shows the canonical amino acid residue in red, the right panel shows the structure with the P34F substitution. d,e) Same as (a), (b) for sg2 screened with ABE. f) Same as (c), but with the E33G substitution. g) Same as (b) and (e). h) Summary of validation results. 1° z-score indicates the average z-scored LFC of the sgRNA in the primary screen. % WT 2° indicates the % of reads that were still WT (unedited) on day 8 of the validation experiment. 2° WT enrichment indicates the average change in the abundance of the WT allele from day 8 to day 21 in the validation experiment. PAM bin is indicated on the left. SV indicates “splice variant”.

This same guide (sg2) was also examined with ABE, and we observed an average of 54.3% editing at positions 4 and 5 of the guide, resulting in an E33G mutant. This allele did not change in abundance across timepoints analyzed (57.9% on day 8, 54.6% day 21), indicating that, in contrast to the P34F mutation introduced by this guide when paired with CBE, the E33G mutation does not have a large effect on cell fitness (**Fig 6d, e**). Indeed, when this mutation was profiled by SGE, it scored as intermediate ^42^. Further, this residue points away from BARD1, and does not come in close contact with the Zn^2+^ atoms or its binding loops (**Fig 6f**).

We also validated sg10 with both base editors. With CBE, we observed the predicted H1767Y mutation, as well as a second Q1768* mutation that resulted from out-of-window editing (26.7% C>T editing at C9). Alleles with the single H1767Y mutation depleted from an average of 35.9% on day 8 to 20.8% of reads on day 21 (**Fig 6g**) while the WT allele enriched from 35.2 to 64.4% of reads. Because alleles containing only the H1767Y mutation were depleted, we conclude that this mutation is sufficient for LOF (**Supplementary Figure 6d**). These results are concordant with the SGE data, as both H1767Y and Q1768* individually score as LOF^42^. With ABE, sg10 introduces either an N1766S mutation (47.2% A>G conversion, A4) or N1766G mutation (18.7% A>G conversion at A3; 47.2% A>G conversion at A4) as well as an H1767R mutation (59.1% A>G conversion at A7), resulting in a LOF phenotype, while the WT allele enriches from 42.1% to 58.7% (**Fig 6g**, **Supplementary Figure 6e**). Given that the H1767R mutation occurred alone and did not deplete substantially (10.5% day 8 to 9.8% day 21), it is likely that the mutants at position 1766 were driving this LOF phenotype. Notably, Findlay et al. classified H1767R as functional, which is concordant with our observation; however, they also found that every missense mutation introduced at N1766 is functional, including N1766S^42^. We did not capture any alleles with a mutation only at this position, so cannot make definitive conclusions about the role of N1766S or N1766G. It remains possible that the observed LOF is due to a combinatorial effect of mutations at N1766 and H1767.

As an additional example, sg9 was screened only with CBE and results in a G1727K mutation (**Supplementary Figure 6f, g**). This residue falls in the BRCT phosphopeptide-binding motif, which is conserved in several DNA-damage repair proteins. It is also responsible for the association of BRCA1 with proteins phosphorylated by ATM, and is implicated in breast and ovarian cancers^44^. Although this exact mutation has not been documented in ClinVar, G1727R and G1727E mutations are pathogenic, and G1727V is categorized as LOF by SGE^42^, indicating that many substitutions are not tolerated at this position. We also screened sg15 with both CBE and ABE, which introduces a C64Y mutation with CBE, and C64R and L63P mutations with ABE (**Supplementary Figure 6h-k)**. By our validation criteria, this guide did not validate with NG-CBE, but did previously validate with WT-Cas9-CBE^33^, and with NG-ABE. While we were unable to parse the effects of these individual mutants based on the spectrum of alleles in our data, C64R scored as LOF with SGE and L63P is pathogenic in ClinVar, so it is likely that both of these mutations contribute to the LOF phenotype.

All validation results are summarized in **Fig 6h**. We identified five guides that introduced deleterious mutations with one or both base editors. Two of these guides mutate the RING domain, and three the BRCT domain, both of which are frequently mutated in tumors. None of the guides that utilized a PAM classified as inactive validated, and sequencing showed little editing at these sites when screened with either CBE or ABE. 3 of 8 guides with intermediate or active PAMs introduced benign edits with both base editors, indicating that these were false positives in the primary screen. In sum, we identified the causal mutations driving the LOF phenotypes in the primary screen for five sgRNAs, all of which utilized an intermediate or active PAM. In contrast, 0 of 10 sgRNAs with inactive PAMs validated, reinforcing the PAM-specificity of these Cas9 variants and highlighting the necessity of validating primary screening results.

#### BCL2

Since its FDA approval in 2016, Venetoclax, which targets the anti-apoptotic protein BCL2, has been administered to thousands of patients with chronic lymphocytic leukemia (CLL), small lymphocytic lymphoma (SLL) or acute myeloid leukemia (AML)^46^. Unfortunately, many patients develop resistance to treatment, causing tumor relapse. Many of these are single amino acid mutations in BCL2 that emerge in patients over the course of treatment, and several other mutations that lead to drug resistance have been characterized in human cells or mice^47–49^. We set out to identify additional resistance-causing mutations in BCL2, as a better understanding of these resistance mechanisms can help improve patient monitoring and allow for tailored treatment plans, as well as inform the design of new, mutation-agnostic drugs.

We designed a tiling library targeting *BCL2* with Cas9-NG, generated both CBE and ABE versions, and screened in triplicate at high coverage (>10,000 cells per sgRNA) in MOLM13 cells, an AML cell line sensitive to BCL2 inhibition. Following selection with puromycin, we treated cells with 62.5 nM Venetoclax for 14 days (**Fig 7a**). LFC values for the untreated cells were calculated relative to plasmid DNA, and LFC values for Venetoclax-treated cells were calculated relative to untreated arms (**Supplementary Data 7**). Correlations between the untreated replicates indicated good technical quality (Pearson’s r = 0.6-0.9) whereas the poor correlation between replicates of the treated arms (Pearson’s r = 0.05-0.55) is attributed to the stringent nature of this positive selection screen, as few guides confer resistance. Next, we calculated z-scores for each guide relative to intergenic controls. 16 guides enriched with a z-score >3 with either or both base editors (**Supplementary Figure 7a**), and 68.8% (11/16) of these are predicted to edit between position 100 and 175, a region that contains the P2 and P4 pockets responsible for binding of Venetoclax (**Fig 7b**). Notably, several sites of resistance mutations observed clinically (G101, D103, F104) fall within this region^47–49^.

**Figure 7.**
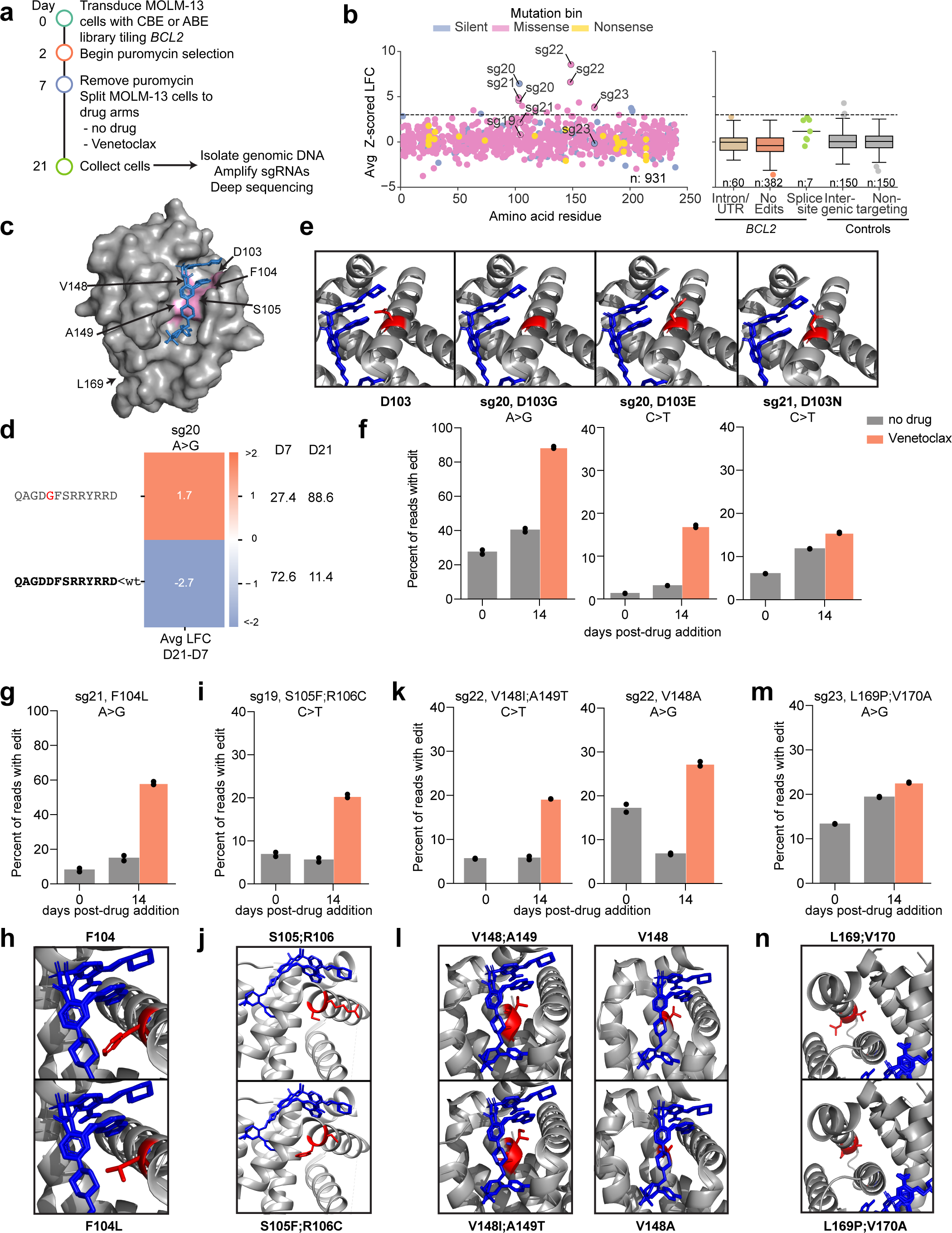
Venetoclax-resistant BCL2 mutants identified by base editing with Cas9-NG. a) Timeline by which tiling screens were conducted. b) Performance of sgRNAs targeting BCL2 for the Venetoclax-treated arm, plotting both CBE and ABE screens, colored according to the predicted mutation bin. A dashed line delineates the z-score cutoff of 3. Boxes show the quartiles; whiskers show 1.5 times the interquartile range. Categories with n < 20 are shown as individual dots. c) 3D structure of BCL2 in complex with Venetoclax (PDB ID: 6O0K). Amino acids that sg19-23 are predicted to edit are highlighted in pink. d) Translated sequence around sg20 for any allele with at least 1% abundance in any condition with ABE. The WT sequence is bolded in black, unchanged amino acids are in grey, and substitutions are highlighted in red. Avg LFC from day 21 - day 7 is indicated on the heatmap and relative percent abundance of each allele is indicated to the right (normalized after filtering for alleles with <1% abundance at both timepoints). e) Structural visualization of WT D103 and the mutations indicated in (f). f) Percentage of reads from the most enriched D103 mutants after 14 days of Venetoclax treatment. sgRNA, edit type, and amino acid mutation are indicated. Dots indicate n=2 biological replicates. g,i,k,m) Same as (f) but for indicated sgRNA, edit type, and position. h,j,l,n) Same as (e) but showing the most enriched mutant indicated in (g),(i),(k), or (m), respectively.

#### BCL2 Validation

In order to confirm the mutations causing these phenotypes, we chose five highly enriched sgRNAs to validate with both base editors, including three guides predicted to make missense edits at residues 103-105 (sg19 - 21), as well as two guides predicted to make missense edits at residues 148/149 (sg22) and 169 (sg23), which are denoted on the 3D structure of BCL2 in complex with Venetoclax (**Fig 7c**). We performed validation screens, as described above for *BRCA1*, with each of the guides individually transduced into MOLM13 cells in duplicate. The conditions from the original *BCL2* tiling screens were replicated, and following isolation and amplification of genomic DNA, we Illumina sequenced the targeted loci.

Upon analysis of the sequencing, we saw levels of editing at the early time point vary from guide to guide, ranging from 11.7 - 55.4% with ABE and 10.9% - 47.9% with CBE in the predicted window (**Supplementary Figure 7b**). While all predicted edits were based on an assumed editing window at positions 4-8, we also observed editing in the 3-10 window with both ABE and CBE, as well as low levels of C>T editing farther afield (sg22) which led to several unpredicted amino acid substitutions (**Supplementary Figure 7b**). As before, we used the relative abundances of the wild type allele at the early and late time points to evaluate whether a guide validated. In this case, if the WT allele depleted by more than 10% under the selective pressure of Venetoclax, we considered it to be validated (**Supplementary Figure 7c**). Based on this criterion, 8 of 8 guide-BE combinations validated as true positives, and 2 of 2 validated as true negatives. Further, sg20, 21, and 22 validated with both ABE and CBE, whereas sg19 and sg23 validated with the base editor with which they scored in the primary screens, but not with the non-scoring base editor; indeed, these sgRNAs were predicted to make either a silent edit (sg23, CBE), or no edit (sg19, ABE). That top hits from the *BCL2* screens had a higher validation rate than those from the *BRCA1* screens is likely because the former is a positive selection screen, which presents fewer opportunities for off-target activity to score as false positives.

We next examined the enrichment of specific alleles to determine the causal resistance mutation(s). For sg20 we observed some C>T editing with CBE at positions 5, 8, and 10 on day 7, but saw strong enrichment for the D103E mutation caused by a C>A transversion at position 5 (**Supplementary Figure 7d**). In this case, all missense mutations were the result of out-of-window edits or transversions, highlighting the necessity for direct sequencing of the edited locus. When paired with ABE, sg20 led to high levels of editing at A4, resulting in strong enrichment for the D103G allele, which likely disrupts the α2 helix (**Fig 7d, e, f**). When we examined sg21 with CBE, we also saw enrichment for missense mutations at D103; substitution for an asparagine (D103N) was most favored upon Venetoclax treatment, but we also observed enrichment of the D103Y allele, as well as the dual replacement of aspartic acids at 102 and 103 with asparagine (**Supplementary Figure 7e**). The D103 residue falls within the P4 pocket and is known to be important for hydrogen binding between the azaindole moiety of Venetoclax and BCL2^48^. Both D103E and D103Y have been previously recorded in patient samples bearing the G101V mutation^48, 50^, and Blombery and colleagues have further shown that D103E mutagenesis causes the P4-binding pocket to more closely resemble that of BCL-XL, which is not inhibited by Venetoclax. When screened with ABE, sg21 predominantly enriched for the F104L mutation that has been shown to increase the P2-binding pocket volume^51^ and likely disrupts a hydrogen bond between Venetoclax and the side chain of F104 (**Fig 7g, h**, **Supplementary Figure 7f**). For sg19 we saw C>T editing at positions 5, 7, 8, and 9, which introduces an S105F missense mutation in 94.5% of edited alleles at the early time point (**Supplementary Figure 7g**). In all cases this mutant enriched during treatment with Venetoclax, and dual editing of S105F and R106C enriched further still (**Fig 7i, j**). Interestingly, the strongest LFC was seen with a rare in-frame deletion that removes R106.

While the preceding three sgRNAs introduced edits at positions 102-106, which are located in the α2 helix, we also saw edits on the α5 helix and the non-core α6 helix enrich in the primary screen. With sg22 we observed the introduction of several resistance-causing missense mutations at positions 148-152. When screened with CBE, an A149T mutation was observed in 94.2% of all edited alleles on day 7 (**Supplementary Figure 7h**), which alone was able to confer resistance, but when A149T occurred in combination with V148I we saw further enrichment (**Fig 7k, l**). When sg22 was screened with ABE, we saw A>G editing at positions 3, 4, 9 and 10, leading to the introduction of a V148A edit in all edited alleles. While this edit alone was sufficient to cause Venetoclax resistance, we also observed secondary edits at position F150 (to L and P) which enriched during drug treatment (**Supplementary Figure 7i**).

The final sgRNA that we validated (sg23) was predicted to edit at position L169. With CBE, sg23 was predicted to make a silent edit, though we did observe low levels of editing in a large window (C0-C18) resulting in non-resistant missense edits at positions V170 and D171. With ABE all edited alleles carried the L169P missense mutation (**Supplementary Figure 7j**).

Interestingly, this edit only enriched when V170A mutagenesis was observed in tandem (**Fig 7m**). This resistance mechanism is particularly interesting, because amino acids 169 and 170 are located on the far side of the protein on the α6 helix and do not come in direct contact with Venetoclax. Mapping of these mutations onto the crystal structure of BCL2 shows the potential of a larger structural impact, whereby substitution with two significantly smaller side chains on the inner face of the helix creates vacated space which may then be compensated for by additional conformational changes in the protein (**Fig 7n**). A final summary of the performance of sg19-23 in both the primary and secondary screens is provided (**Supplementary Figure 7k**).

By leveraging PAM-flexible Cas9-NG, and both A-G and C-T base editors, we were able to densely tile *BCL2* and identify nucleotide substitutions that confer resistance to Venetoclax. Analysis of the edited sequences obtained from Venetoclax-treated samples confirmed three previously-documented mutations (F104L, D103E, D103Y), and revealed several resistant mutations that, to our knowledge, have not been reported. This screen demonstrates the power of tiling base editing screens in a positive selection setting, and identifies a condensed region of *BCL2* (100-175) harboring many resistance mutations, which may be of particular interest for additional experiments with more exhaustive forms of mutagenesis.

## DISCUSSION

We have established a pipeline that allows for the profiling of new Cas variants and applied it to high-fidelity variants generated to mitigate off-target effects as well as PAM-flexible variants that increase the targeting range of Cas9. With respect to the former, we find that WT-Cas9 is still the best option for most screening applications because it does not require the use of G19 guides, and thus there are approximately four-fold more guides available. However, if off-target effects are of a particular concern, then eCas9-1.1 is the best option of those tested here. It has recently been shown that so-called “Blackjack mutations” improve the on-target activity of these high fidelity variants with 5’G extended sgRNAs^28^, and one such nuclease, eSpCas9-plus could be useful in mitigating off-target effects without compromising guide selection options. Importantly, here we benchmark these variants in the context of genetic screens in which both the Cas9 and guide are delivered via lentivirus, and thus must complex together intracellularly. In cases where the Cas9 protein is complexed with the guide *in vitro*, i.e. RNP delivery, the binding conditions are extraordinarily more favorable, which may prevent much of the on-target activity loss observed in our benchmarking for high-fidelity variants that have performed well when delivered as an RNP.

We also screened five PAM-flexible variants, two of which showed promising activity at non-canonical PAM sequences. We directly compared the off-target profiles of Cas9-NG and SpG, and found that they are nearly indistinguishable. Given that SpG shows higher activity at several more PAM sites than Cas9-NG, with a comparable off-target profile, we recommend performing base editing screens with this enzyme going forward. Indeed, due to the relatively high prevalence of false-negatives with the technology, especially in negative-selection screens, the benefit of added depth is worthwhile, enabling the use of multiple unique guides to pinpoint regions of particular interest in a target locus. The recent development of a nearly-PAM-less Cas9 variant^26^, as well as approaches to generate C>G edits^52–55^, suggests that more editing outcomes and thus even finer resolution will be possible for base editor screens. Generating a library of every possible amino acid substitution for an entire open reading frame is certainly possible^56–58^, but is also expensive. By highlighting specific protein regions of high value, densely tiled base editing screens can thus guide the creation of smaller, more-focused ORF libraries, or SGE approaches, that are commensurately more efficient to screen. The recent demonstration of prime editing technology for focused, saturating mutagenesis on haploidized loci^59^ provides another potential path for follow-up of base editing screens.

Positive selection screens are generally easier to execute with a lower false positive rate compared to negative selection screens^60^, a trend consistent with the validation rate of the *BCL2* and *BRCA1* screens presented here, as well as our previously successful drug resistance screens for inhibitors of *MCL1, BCL2L1,* and *PARP1*^33^. The low upfront costs of generating a pooled, base editing library, coupled with the small-scale and relative ease of execution, suggests that such screens can be used to, for example, identify a resistance mutation that helps to prove the actual target of a less-characterized small molecule, as well as gain insight into potential resistance mechanisms long before seeing what arises in patients. As cell line models are likely to already exist for probing the activity of such small molecules, there are few barriers to implementing such screens early in the drug discovery process.

## ACKNOWLEDGEMENTS

We thank Amy Goodale, Briana Fritchman, Edith Sawyer, Hinako Kiwabe, Luke Sprenkle, Pema Tenzing, Tashi Lokyitsang and Xiaoping Yang for producing guide libraries and lentivirus; Olivia Bare, Yenarae Lee, and Max Macaluso for logistics support; Matthew Greene, Bronte Wen, Adam Brown, Doug Alan, Mark Tomko, and Tom Green for software engineering support; the Broad Institute Genomics Platform Walk-up Sequencing group for Illumina sequencing; and the Functional Genomics Consortium for funding support. We thank Marissa Feeley, Sarah Weiss, Nicky Persky, Dave Root, Kathleen Kristie, Russell Walton, Sumaiya Iqbal, and Benjamin Kleinstiver for helpful discussions.

## AUTHOR CONTRIBUTIONS

Conceived of the study: AKS, JGD

Executed genetic screens: AKS, ALG, ZMS, AVM, REH

Performed analyses: AKS, ALG, PR, ZMS, PCD, MH

Created visualizations: AKS, ALG, ZMS, PCD

Designed libraries: AKS, MH, ALG

Curated data: AKS, ALG, ZMS, MH

Wrote the manuscript: AKS, ALG, JGD

Supervised the project: JGD

## COMPETING INTERESTS

JGD consults for Microsoft Research, Agios, Phenomic AI, Maze Therapeutics, BioNTech, and Pfizer; JGD consults for and has equity in Tango Therapeutics. JGD’s interests were reviewed and are managed by the Broad Institute in accordance with its conflict of interest policies. All other authors declare no competing interests.

## METHODS

### Vectors

pRosetta (Addgene 59700): lentiviral construct for expression of eGFP, puromycin resistance and blasticidin resistance.

pRosetta_v2 (Addgene 136477): modification of pRosetta to include a hygromycin resistance cassette; also known as pRDA_018.

pRDA_118 (Addgene 133459): U6 promoter expresses customizable SpCas9 guide; EF1a promoter provides puromycin resistance. This vector is a derivative of the lentiGuide vector, with a modification to the tracrRNA to eliminate a run of four thymidines.

pRDA_091: U6 promoter expresses customizable SpCas9 guide; EF1a promoter provides puromycin resistance. This vector also contains a Tet3G cassette that was not utilized in this study.

All Cas9 variants have: EF1a expresses Cas9; T2A site provides blasticidin resistance and P2A site provides mKate2. The Cas9 variants were generated by introducing the point mutations (Genscript) described in the original publications.

pRDA_085 (Addgene 158583): WT-Cas9.

pRDA_151 (Addgene TBD) : Cas9-HF1^9^. Point mutations: N497A/ R661A/ Q695A/ Q926A

pRDA_152 (Addgene TBD) : eCas9-1.1^11^. Point mutations: K848A/ K1003A/ R1060A

pRDA_153 (Addgene TBD) : evoCas9^13^. Point mutations: M495V/ Y515N/ K526E/ R661Q

pRDA_154 (Addgene TBD) : xCas9-3.7^23^. Point mutations: A262T/ R324L/ S409I/ E480K/ E543D/ M694I/ E1219V

pRDA_155 (Addgene TBD) : Cas9-VQR^21^. Point mutations: D1135V /R1335Q/ T1337R

pRDA_156 (Addgene TBD) : Cas9-VRER^21^. Point mutations: D1135V /G1218R/ R1335E/ T1337R

pRDA_157 (Addgene TBD) : HypaCas9^10^. Point mutations: N692A/ M694A/ Q695A/ H698A

pRDA_275 (Addgene TBD) : Cas9-NG^22^. Point mutations: L1111R/ D1135V/ G1218R/ E1219F/ A1322R/ R1335V/ T1337R

pRDA_381 (Addgene TBD) : HiFi Cas9^12^. Point mutations: R691A

pRDA_449 (Addgene TBD) : SpG^26^. Point mutations: D1135L/ S1136W/ G1218K/ E1219Q/ R1335Q/ T1337R

All base editor constructs have: U6 promoter expresses customizable guide RNA with a 10x guide capture sequence at the 3’ end of the tracrRNA to facilitate future use with direct capture single cell RNA sequencing^61^; core EF1a (EFS) expresses codon-optimized ABE or CBE, and 2A site provides puromycin resistance. Note that the ABE8e constructs contain the V106W mutation.

pRDA_256 (Addgene 158581): WT-BE3

pRDA_336 (Addgene TBD): NG-BE3

pRDA_478 (Addgene TBD): SpG-BE3

pRDA_426 (Addgene TBD): WT-ABE8e pRDA_429 (Addgene TBD): NG-ABE8e

pRDA_479 (Addgene TBD): SpG-ABE8e

### Cell lines and culture

A375, MOLM13 and MELJUSO cells were obtained from Cancer Cell Line Encyclopedia at the Broad Institute. Unmodified HAP1 cells (item C631) were obtained from Horizon Discovery. HEK293Ts were obtained from ATCC (CRL-3216). MOLM13 cells were selected based on data from genome-wide CRISPR screens and cancer cell drug sensitivity screens (CTD^2 and GDSC) found on the Cancer Dependency Map Portal which identified MOLM13 as dependent on *BCL2*.

All cells regularly tested negative for mycoplasma contamination and were maintained in the absence of antibiotics except during screens, validation experiments, and lentivirus production, during which media was supplemented with 1% penicillin-streptomycin. Cells were passaged every 2-4 days to maintain exponential growth and were kept in a humidity-controlled 37°C incubator with 5.0% CO2. Media conditions and doses of polybrene, puromycin, blasticidin, and hygromycin were as follows, unless otherwise noted:

A375: RPMI + 10% fetal bovine serum (FBS); 1 µg/mL; 1 µg/mL; 5 µg/mL; N/A

HAP1: IMDM + 10% FBS; 4 µg/mL; 2 µg/mL; 5 µg/mL; N/A

HEK293T: DMEM + 10% heat-inactivated FBS; N/A; N/A; N/A; N/A

MELJUSO: RPMI + 10% FBS; 4 µg/mL; 1 µg/mL; 4 µg/mL; 100 µg/mL

MOLM13: RPMI + 10% FBS; 4 µg/mL; 1 µg/mL; N/A; N/A

### PAM-mapping library design

50 essential and non-essential genes were picked from prior screens performed in A375 and HT29. *BRCA1* and *BRCA2* were also included to increase the coverage per PAM sequence. sgRNA sequences tiling the coding sequence of the principal Ensembl transcript of these genes were designed. Four nucleotides following the sgRNA sequence were reported as the PAM sequence. The library was filtered to exclude any sgRNAs with BsmBI sites or a TTTT sequence. Promiscuous guides (defined as the PAM-proximal 18mer having >=5 off-targets with up to 1 mismatch in the genome) were filtered out. We aimed to pick 50 sgRNAs per PAM sequence for the essential genes, *BRCA1,* and *BRCA2* and 25 sgRNAs per PAM sequence for non-essential genes. In doing so, we picked 10 G19, 30 g20 and 10 G20 sgRNAs for essential genes and 5 G19, 15 g20 and 5 G20 sgRNAs for non-essential genes.

### *BRCA1* base editor tiling library design

Guide sequences for tiling libraries were designed using sequence annotations from Ensembl (GRCg38). We used Ensembl’s REST API (https://rest.ensembl.org/) to obtain the genomic locations of transcripts, transcript sequences, and protein sequences, and used these to annotate each sgRNA with its predicted edits. We included all sgRNAs targeting the coding sequence; we also included all sgRNAs for which the start was up to 30 nucleotides into the intron and UTRs. We designed every possible guide (using an NNNN PAM) against the longest annotated transcript for *BRCA1* (ENST00000471181, 1884 amino acids) using an editing window of 4-8 nucleotides for both CBE and ABE screens. We filtered out guides >5 perfect matches in the genome. The library was filtered to exclude any sgRNAs with BsmBI sites or a TTTT sequence.

### *BCL2* base editor tiling library design

We used the Ensembl transcript ENST00000333681.5 to design all guides targeting *BCL2*, regardless of PAM, annotating edits based on an editing window of 4-8; we also included all sgRNAs for which the start was up to 29 nucleotides into the intron and UTRs. Guides with PAMs that scored as a fraction active s; 0.1 (from the PAM tiling screen) were filtered out for a total of n = 96 inactive PAMs in the library. Finally, the library was filtered to exclude any sgRNAs with BsmBI sites or a TTTT sequence.

### Library production

Oligonucleotide pools were synthesized by CustomArray. BsmBI recognition sites were appended to each sgRNA sequence along with the appropriate overhang sequences (bold italic) for cloning into the sgRNA expression plasmids, as well as primer sites to allow differential amplification of subsets from the same synthesis pool. The final oligonucleotide sequence was thus: 5’-[Forward Primer]CGTCTCA***CACCG***[sgRNA, 20 nt]***GTTT***CGAGACG[Reverse Primer].

Primers were used to amplify individual subpools using 25 µL 2x NEBnext PCR master mix (New England Biolabs), 2 µL of oligonucleotide pool (-40 ng), 5 µL of primer mix at a final concentration of 0.5 µM, and 18 µL water. PCR cycling conditions: (1) 98°C for 30 seconds; (2) 53°C for 30 seconds; (3) 72°C for 30 seconds; (4) go to (1), x 24.

In cases where a library was divided into subsets, unique primers could be used for amplification:

Primer Set; Forward Primer, 5’ - 3’; Reverse Primer, 5’ - 3’

1; AGGCACTTGCTCGTACGACG; ATGTGGGCCCGGCACCTTAA

2; GTGTAACCCGTAGGGCACCT; GTCGAGAGCAGTCCTTCGAC

3; CAGCGCCAATGGGCTTTCGA; AGCCGCTTAAGAGCCTGTCG

4; CTACAGGTACCGGTCCTGAG; GTACCTAGCGTGACGATCCG

5; CATGTTGCCCTGAGGCACAG; CCGTTAGGTCCCGAAAGGCT

6; GGTCGTCGCATCACAATGCG; TCTCGAGCGCCAATGTGACG

The resulting amplicons were PCR-purified (Qiagen) and cloned into the library vector via Golden Gate cloning with Esp3I (Fisher Scientific) and T7 ligase (Epizyme); the library vector was pre-digested with BsmBI (New England Biolabs). The ligation product was isopropanol precipitated and electroporated into Stbl4 electrocompetent cells (Invitrogen) and grown at 30°C for 16 h on agar with 100 µg/mL carbenicillin. Colonies were scraped and plasmid DNA (pDNA) was prepared (HiSpeed Plasmid Maxi, Qiagen). To confirm library representation and distribution, the pDNA was sequenced.

### Lentivirus production

For small-scale virus production, the following procedure was used: 24 h before transfection, HEK293T cells were seeded in 6-well dishes at a density of 1.5 x 10^6^ cells per well in 2 mL of DMEM + 10% heat-inactivated FBS. Transfection was performed using TransIT-LT1 (Mirus) transfection reagent according to the manufacturer’s protocol. Briefly, one solution of Opti-MEM (Corning, 66.75 µL) and LT1 (8.25 µL) was combined with a DNA mixture of the packaging plasmid pCMV_VSVG (Addgene 8454, 250 ng), psPAX2 (Addgene 12260, 1250 ng)^62^, and the transfer vector (e.g., pLentiGuide, 1250 ng). The solutions were incubated at room temperature for 20-30 min, during which time media was changed on the HEK293T cells. After this incubation, the transfection mixture was added dropwise to the surface of the HEK293T cells, and the plates were centrifuged at 1000 g for 30 min at room temperature. Following centrifugation, plates were transferred to a 37°C incubator for 6-8 h, after which the media was removed and replaced with DMEM +10% FBS media supplemented with 1% BSA. Virus was harvested 36 h after this media change.

A larger-scale procedure was used for pooled library production. 24 h before transfection, 18 x 10^6^ HEK293T cells were seeded in a 175 cm^2^ tissue culture flask and the transfection was performed the same as for small-scale production using 6 mL of Opti-MEM, 305 µL of LT1, and a DNA mixture of pCMV_VSVG (5 µg), psPAX2 (50 µg), and 40 µg of the transfer vector. Flasks were transferred to a 37°C incubator for 6-8 h; after this, the media was aspirated and replaced with BSA-supplemented media. Virus was harvested 36 h after this media change.

### Determination of antibiotic dose

In order to determine an appropriate antibiotic dose for each cell line, cells were transduced with the pRosetta or pRosetta_v2 lentivirus such that approximately 30% of cells were transduced and therefore EGFP+. At least 1 day post-transduction, cells were seeded into 6-well dishes at a range of antibiotic doses (e.g. from 0 µg/mL to 8 µg/mL of puromycin). The rate of antibiotic selection at each dose was then monitored by performing flow cytometry for EGFP+ cells. For each cell line, the antibiotic dose was chosen to be the lowest dose that led to at least 95%

EGFP+ cells after antibiotic treatment for 7 days (for puromycin) or 14 days (for blasticidin and hygromycin).

### Small molecule doses in pooled screens

For *BRCA1* screens in MELJUSO cells, cisplatin (BioVision, 1550) was diluted in 0.9% NaCl and was screened at 1 µM. For *BCL2* screens in MOLM13 cells, Venetoclax (Selleckchem, S8048) was diluted in DMSO and was screened at 62.5 nM.

### Determination of lentiviral titer

To determine lentiviral titer for transductions, cell lines were transduced in 12-well plates with a range of virus volumes (e.g. 0, 150, 300, 500, and 800 µL virus) with 1 to 3 x 10^6^ cells per well in the presence of polybrene. The plates were centrifuged at 640 x g for 2 h and were then transferred to a 37°C incubator for 4-6 h. Each well was then trypsinized, and an equal number of cells seeded into each of two wells of a 6-well dish. Two days post-transduction, puromycin was added to one well out of the pair. After 5 days, both wells were counted for viability. A viral dose resulting in 30-50% transduction efficiency, corresponding to an MOI of -0.35-0.70, was used for subsequent library screening.

### Derivation of stable cell lines

In order to establish Cas9 variant expressing cell lines for screens with the PAM-mapping tiling library and both off-target libraries, A375 cells were transduced with either pRDA_085, pRDA_151-157, pRDA_275, pRDA_381 or pRDA_449 and successfully transduced cells were selected with blasticidin for a minimum of 2 weeks. Cells were taken off blasticidin at least one passage before transduction with libraries.

### Pooled screens

For pooled screens, cells were transduced in 2-3 biological replicates with the lentiviral library. Transductions were performed at a low multiplicity of infection (MOI ∼0.5), using enough cells to achieve a representation of at least 500 transduced cells per sgRNA assuming a 20-40% transduction efficiency. For the CRISPRko screens, cells were plated in polybrene-containing media with 3 x 10^6^ cells per well in a 12-well plate. Because the titer of all-in-one base editor viruses was low, cells were plated in polybrene-containing media with 1.5 x 10^6^ cells per well in a 12-well plate. Plates were centrifuged for 2 hours at 640 x g, after which 2 mL of media was added to each well. Plates were then transferred to an incubator for 4-6 hours, after which virus-containing media was removed and cells were pooled into flasks. Puromycin was added 2 days post-transduction and maintained for 5-7 days to ensure complete removal of non-transduced cells. Upon puromycin removal, cells were split to any drug arms (each at a representation of at least 1,000 cells per sgRNA) and passaged every 2-4 days for an additional 2 weeks to allow sgRNAs to enrich or deplete; cell counts were taken at each passage to monitor growth.

### Genomic DNA isolation and sequencing

Genomic DNA (gDNA) was isolated using the KingFisher Flex Purification System with the Mag-Bind® Blood & Tissue DNA HDQ Kit (Omega Bio-Tek). The gDNA concentrations were quantitated by Qubit. For samples where genomic DNA was limiting, gDNA was purified prior to PCR using the Zymo OneStep PCR Inhibitor Removal Kit (Zymo), per the manufacturer’s instructions.

For PCR amplification, gDNA was divided into 100 µL reactions such that each well had at most 10 µg of gDNA. Plasmid DNA (pDNA) was also included at a maximum of 100 pg per well. Per 96-well plate, a master mix consisted of 150 µL DNA Polymerase (Titanium Taq; Takara), 1 mL of 10x buffer, 800 µL of dNTPs (Takara), 50 µL of P5 stagger primer mix (stock at 100 µM concentration), 500 µL of DMSO (if used), and water to bring the final volume to 4 mL. Each well consisted of 50 µL gDNA and water, 40 µL PCR master mix, and 10 µL of a uniquely barcoded P7 primer (stock at 5 µM concentration). PCR cycling conditions were as follows: (1) 95°C for 1 minute; (2) 94°C for 30 seconds; (3) 52.5°C for 30 seconds; (4) 72°C for 30 seconds; (5) go to (2), x 27; (6) 72°C for 10 minutes. PCR primers were synthesized at Integrated DNA Technologies (IDT). PCR products were purified with Agencourt AMPure XP SPRI beads according to manufacturer’s instructions (Beckman Coulter, A63880), using a 1:1 ratio of beads to PCR product. Samples were sequenced on a HiSeq2500 HighOutput (Illumina) with a 5% spike-in of PhiX.

### Validation experiments

For validation experiments in which the target site was directly sequenced, individual sgRNAs were cloned into either pRDA_336 (NG-CBE) or pRDA_429 (NG-ABE) and made into lentivirus as described above. At least 1.5 x 10^6^ cells were transduced in duplicate with a virus volume to obtain -30-50% transduction efficiency and were selected with puromycin for 5-7 days to remove untransduced cells; puromycin doses were as described above. After puromycin selection was removed, cells were split into any drug arms and cultured for an additional 14 days. Cell pellets were collected on days 7, 14, and 21 (*BCL2)* or 8, 14, and 21 (*BRCA1*).

Genomic DNA was isolated using either the Kingfisher as described above, or cells were lysed in 96-well plates using 25 µL per well of Lucigen QuickExtract DNA Extraction Solution (QE0905T). Briefly, 25uL of lysis buffer was added to each well, plate was sealed and vortexed, then heated at 65°C for 15 minutes, heated at 95°C for 5 minutes and then stored at -20°C. Target sites were amplified using a 2-step PCR. For the samples in which gDNA was isolated using the Kingfisher, in the first round of PCR, genomic DNA was amplified using custom primers designed to amplify each target site (see **Supplementary Data 8**). Each well contained 50 µL of NEBNext High Fidelity 2X PCR Master Mix (New England Biolabs), 0.5 µL of each primer at 100 µM, and 49 µL of gDNA. We used a touchdown PCR with the following cycling conditions: (1) 98°C for 1 minute; (2) 98°C for 30 seconds; (3) 68°C for 30 seconds (- 1° per cycle); (4) 72°C for 1 minute; (5) Go to step 2, x 15; (6) 72°C for 10 minutes. For samples subjected to the 96-well plate lysis, in the first step, the master mix for each 96-well plate consisted of: 75 µL Titanium Taq polymerase, 500 µL 10X Titanium Taq buffer, 400 µL dNTPs, 250 µL DMSO, 25 µL forward primer at 100 µM, 25 µL reverse primer at 100 µM, and water to bring the final volume to 10 mL. Forward and reverse primers were as described in **Supplementary Data 8**. Each well consisted of 5 µL crude lysate, 25.5 µL master mix, and water to 50 µL final volume. PCR cycling conditions were as follows: (1) 95°C for 5:00, (2) 94°C for 0:30, (3) 53°C for 0:30, (4) 72°C for 0:20, (5) Go to step 2, x 17 cycles, (6) 72°C for 10:00.

The second round of PCR was the same for both approaches. It appended Illumina adapters and well barcodes for sequencing using the P5 primer “Argon” and the P7 primer “Kermit”. Each well contained 1.5 µL of Titanium Taq (Takara), 10 µL of Titanium Taq buffer, 8 µL of dNTPs, 5 µL of DMSO, 0.5 µL of P5 primer at 100 µM, 10 µL of P7 primer, 55 µL of water, and 10 µL of PCR product from the first PCR. The following cycling conditions were used: (1) 95°C for 1 minute; (2) 94°C for 30 seconds; (3) 52.5°C for 30 seconds; (4) 72°C for 30 seconds; (5) go to (2), x 15; (6) 72°C for 10 minutes. Samples were pooled and purified by primer pair with Agencourt AMPure XP SPRI beads according to the manufacturer’s instructions (Beckman Coulter, A63880), using a 1:1 ratio of beads to PCR product. DNA concentration was quantified using a Qubit and purified samples were pooled proportionally to their concentrations. The pooled library was quantified by Qubit and sequenced using the Illumina MiSeq with a 300 nucleotide single read and a 10% PhiX spike-in.

## QUANTIFICATION AND STATISTICAL ANALYSIS

### Screen analysis

Guide sequences were extracted from sequencing reads by running the PoolQ tool with the search prefix “CACCG” (https://portals.broadinstitute.org/gpp/public/software/poolq). Reads were counted by alignment to a reference file of all possible guide RNAs present in the library. The read was then assigned to a condition (e.g. a well on the PCR plate) on the basis of the 8 nt index included in the P7 primer. Following deconvolution, the resulting matrix of read counts was first normalized to reads per million within each condition by the following formula: read per guide RNA / total reads per condition x 1e6. Reads per million was then log2-transformed by first adding one to all values, which is necessary in order to take the log of guides with zero reads.

Prior to further analysis, we filtered out sgRNAs for which the log-normalized reads per million of the pDNA was > 3 standard deviations from the mean. We also filtered out any sgRNAs containing more than 5 off-target sites in the human genome (for non-NGGN guides) or containing more than 5 off-target sites in the human genome with a CFD score of 1.0 (indicating a perfect or near-perfect match) for NGGN guides. We then calculated the log2-fold-change between conditions. All dropout (no drug) conditions were compared to the plasmid DNA (pDNA); drug-treated conditions were compared to the time-matched dropout sample, with the exception of MELJUSO cells in the *BRCA1* screens, which were compared to the plasmid DNA because loss of *BRCA1* had some viability effect in the absence of drug. We assessed the correlation between log2-fold-change values of replicates.

### Analysis of PAM-mapping screens

The initial PAM characterization screens were carried out in four rounds, each sequenced separately. pDNA raw reads were summed across screens, and log-normalized. Because *BRCA1* and *BRCA2* are not widely panlethal, guides targeting these genes were excluded from all analyses downstream of the calculation of log2-fold-changes. Precision-recall was calculated using sgRNAs targeting essential genes as positive controls, and nonessential genes as negative controls. Fraction active was calculated by quantifying the fraction of guides targeting essential genes that were more depleted than the 5th percentile of the most active nonessential guides with the same PAM, and all non-targeting guides. ROC-AUC was calculated using guides targeting essential genes as positive controls, and guides targeting nonessential genes as negative controls.

### Off-target analyses

Using the perfect match guides as true positives and the 1000 control guides as true negatives, we fit a logistic regression for each condition to predict whether each guide is a perfect match or control based on its log2-fold-change. We then used the fit model to map from log2-fold-changes to the probability of being active (i.e. being a perfect match sgRNA). A value near 1 indicates a guide is active and a value near 0 indicates a guide is inactive. These values were then used to calculate the CFD scores for all enzymes screened (**provided in Supplementary Data 4**). For example, if the interaction between the sgRNA and DNA has a single rG:dA mismatch in position 6, then that interaction receives a score of 0.67. If there are two or more mismatches, then individual mismatch values are multiplied together. For example, an rG:dA mismatch at position 7 coupled with an rC:dT mismatch at position 10 receives a CFD score of 0.57 x 0.87 = 0.50. For the high fidelity off-target screens, these data were also used to predict the activity at double mismatches. Using the same logistic regression model as described above, we defined any double mismatch guide with a > 50% probability of being as active as a perfect match based on its log2-fold-change as active. We used this cutoff to define true positives and ranked by the multiplied single mismatch probabilities to predict the activity of the double mismatch guides. For the off-target screens with Cas9-NG and SpG, data were first filtered for PAMs that were intermediate/active with either enzyme and the top 18 perfect match guides (even if they didn’t have an intermediate/active PAM) were maintained.

### Base editing analyses

To obtain a “mutation bin” for each sgRNA, we ordered the mutation types as: Nonsense > Splice site > Missense > Intron > Silent > UTR. Guides containing multiple mutation types were binned as the most severe mutation type. Guides predicted to make no edits in the editing window were binned as “No edits”. To obtain a “clinical significance bin,” (for *BRCA1* only) we classified sgRNAs predicted to introduce multiple ClinVar SNPs based on the most severe clinical significance: Pathogenic > Likely pathogenic; Pathogenic / Likely pathogenic > Uncertain significance > Conflicting reports of pathogenicity > Variant not listed in ClinVar > Likely benign; Benign / Likely benign > Benign. With this ordering, sgRNAs were only binned as “Likely benign” or “Benign” if they did not introduce any mutations not listed in ClinVar, which effectively have an unknown functional significance.

For the calculations regarding the number of targetable residues in *BRCA1* shown in **Fig 5a**, we considered guides included in the libraries we screened with (filtered for off-targets, BsmBI sites and 4Ts targeting *BRCA1* n = 11,524) with PAMs that were intermediate/active with variant base editors or active (NGGN) for WT base editors. For the combination of ABE and CBE, we took the unique set of the sum of residues editable with either CBE or ABE.

### Analysis of deep sequencing data from validation experiments

CRISPResso2 (version 2.0.30) was used to process all sequencing reads from validation experiments^63^. CRISPResso2 was run in base editor mode using the default settings with the following changes: --min_average_read_quality 25, --w 20. We also set custom values for each sgRNA for --wc, --exclude_bp_from_left, --exclude_bp_from_right, and --default_min_aln_score; these parameters can be found in **Supplementary Data 8**.

To calculate replicate correlations, we used the “Alleles_frequency_table_around_sgRNA” file from the CRISPResso2 output, which contains the read counts for each allele (defined as a subsequence around the sgRNA). We then log-normalized the read counts for each sample (using the same formula described in the “Screen analysis” section). Finally, we filtered out any alleles with < 100 reads in all replicates and drug conditions for that sgRNA, and calculated the Pearson correlation between log-normalized reads.

For further analysis of alleles, we set a more stringent filter in order to avoid spurious log2-fold-change values due to low read counts. We filtered out any alleles that comprised < 1% of the total reads in all replicates and drug conditions for that sgRNA.

### External datasets

The data used for the comparison to our previously generated WT-Cas9 CFD shown in **Supplementary Figure 2b** were obtained from the original publication^6^. For the comparison to Legut et al., we obtained data from the original publication^24^ and considered the CD45 and CD55 data, excluding guides targeting the promoter region. We calculated log2-fold-changes by subtracting the bottom_bin from the top_bin for each Cas, z-scored these data based on the non-targeting controls, then calculated the median z-score of each 3 nucleotide PAM for both this, and our dataset. The data used for the comparison to Kim et al. were obtained from the original publication^31^. We used the average indel frequencies at target sequences grouped by 4-nucleotide PAM in the comparisons shown in **Supplementary Figure 3a-c**. Saturation genome editing data was accessed from the original publication^42^. Because the libraries described in our study and the SGE study were designed against different transcripts (SGE used ENST00000357654), we converted the amino acid positions in the SGE data to be consistent with the transcript we designed against for the comparisons between our validation data and this gold standard dataset.

### Data visualization

Figures were created with Python3 and GraphPad Prism (version 8). Schematics were created with BioRender.com. PyMOL (version 2.3.2) was used to map the screening data onto the following crystal structures from the Protein Data Bank: IJM7 (BRCA1/BARD1 RING-domain heterodimer) and 6O0K (BCL2 bound to Venetoclax). The following commands were used to visualize the BRCA1/BARD1 RING-domain heterodimer in PyMOL (from Walton et al. 2020):

cmd.set(“bg_rgb”, ’white’) cmd.set(“ambient”, ’0.21000’) cmd.set(“direct“, ‘0.40000’)

cmd.set(“reflect”, ‘0.43000’) cmd.set(“power”, ‘2.00000’) cmd.set(“spec_reflect”, ‘-0.01000’)

cmd.set(“line_width”, ‘3.00000’) cmd.set(“cache_display”, ‘off’) cmd.set(“shininess”, ‘30.00000’)

cmd.set(“cartoon_sampling”, ‘7’) cmd.set(“cartoon_loop_radius”, ‘0.15000’)

cmd.set(“cartoon_oval_length”, ‘1.00000’) cmd.set(“auto_color_next”, ‘1’)

cmd.set(“max_threads”, ‘4’) cmd.set(“specular_intensity”, ‘0.30000’) cmd.set(“button_mode_name”, ‘3-Button Viewing’) cmd.set(“seq_view”, ‘on’) cmd.set(“cartoon_ring_mode”, ‘3’)

## DATA AVAILABILITY

The read counts for all screening data and subsequent analyses are provided as Supplementary Data. Fastq files are deposited with the Gene Expression Omnibus (GSE180351) and the Sequence Read Archive (PRJNA753064).

## STATISTICAL ANALYSIS

All z-scores and Pearson correlation coefficients were calculated in Python.

## CODE AVAILABILITY

All custom code used for analysis and example notebooks are available on GitHub: https://github.com/gpp-rnd/cas9-variants-manuscript.

## Supplementary Figures

**Supplementary Figure 1:**
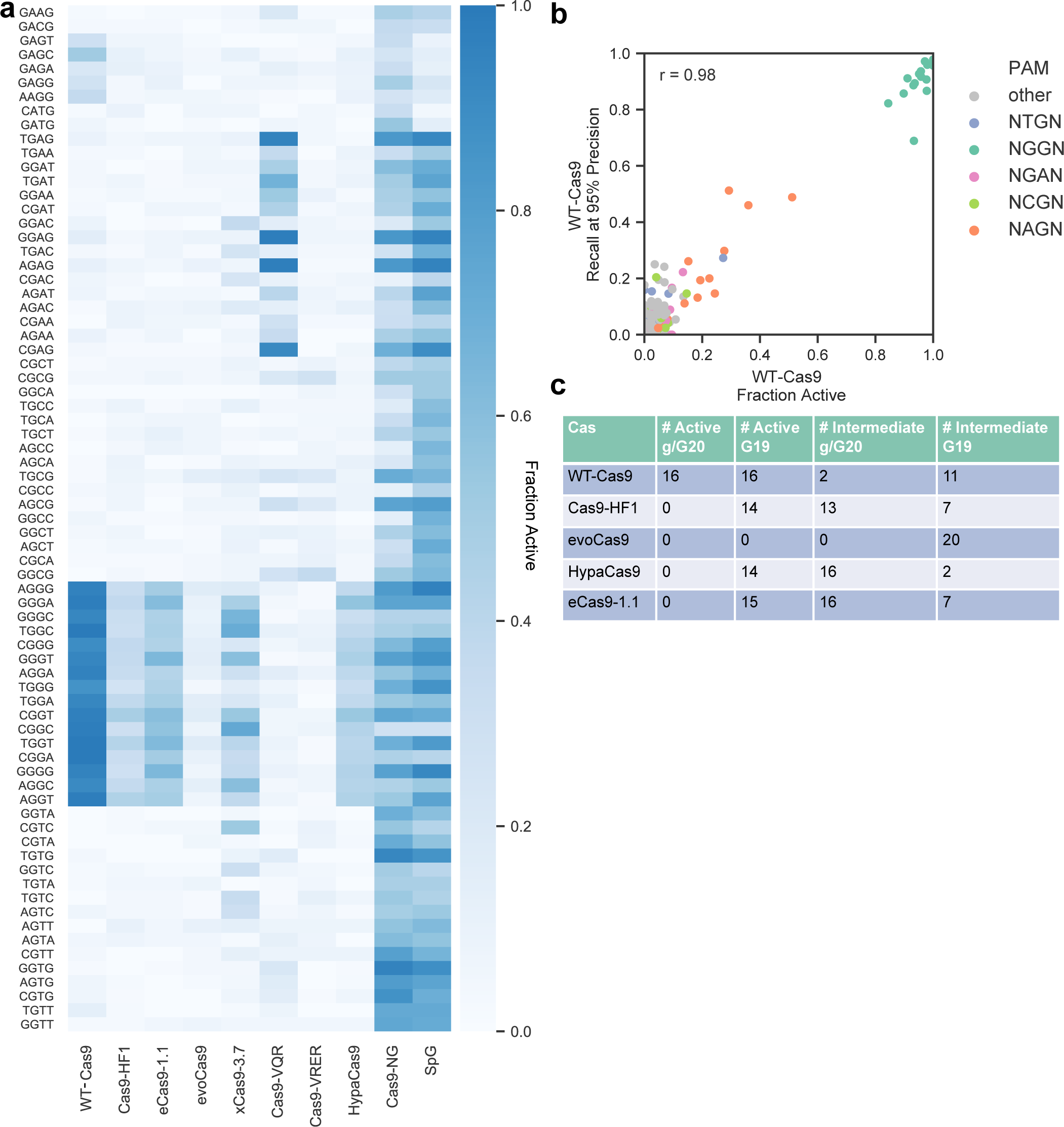
Establishment of a benchmarking assay for Cas9 activity. a) Heatmap of fraction active values for each variant. PAMs that have a fraction active of >= 0.3 with at least one variant are shown on the y-axis. b) Comparison of fraction active metric (x-axis) and recall at 95% precision metric (y-axis) applied to WT-Cas9. Each dot represents a 4 nucleotide PAM, shaded according to the legend on the right. c) Number of active and intermediate PAMs when considering G19 or g/G20 sgRNAs for WT-Cas9 and high fidelity variants.

**Supplementary Figure 2:**
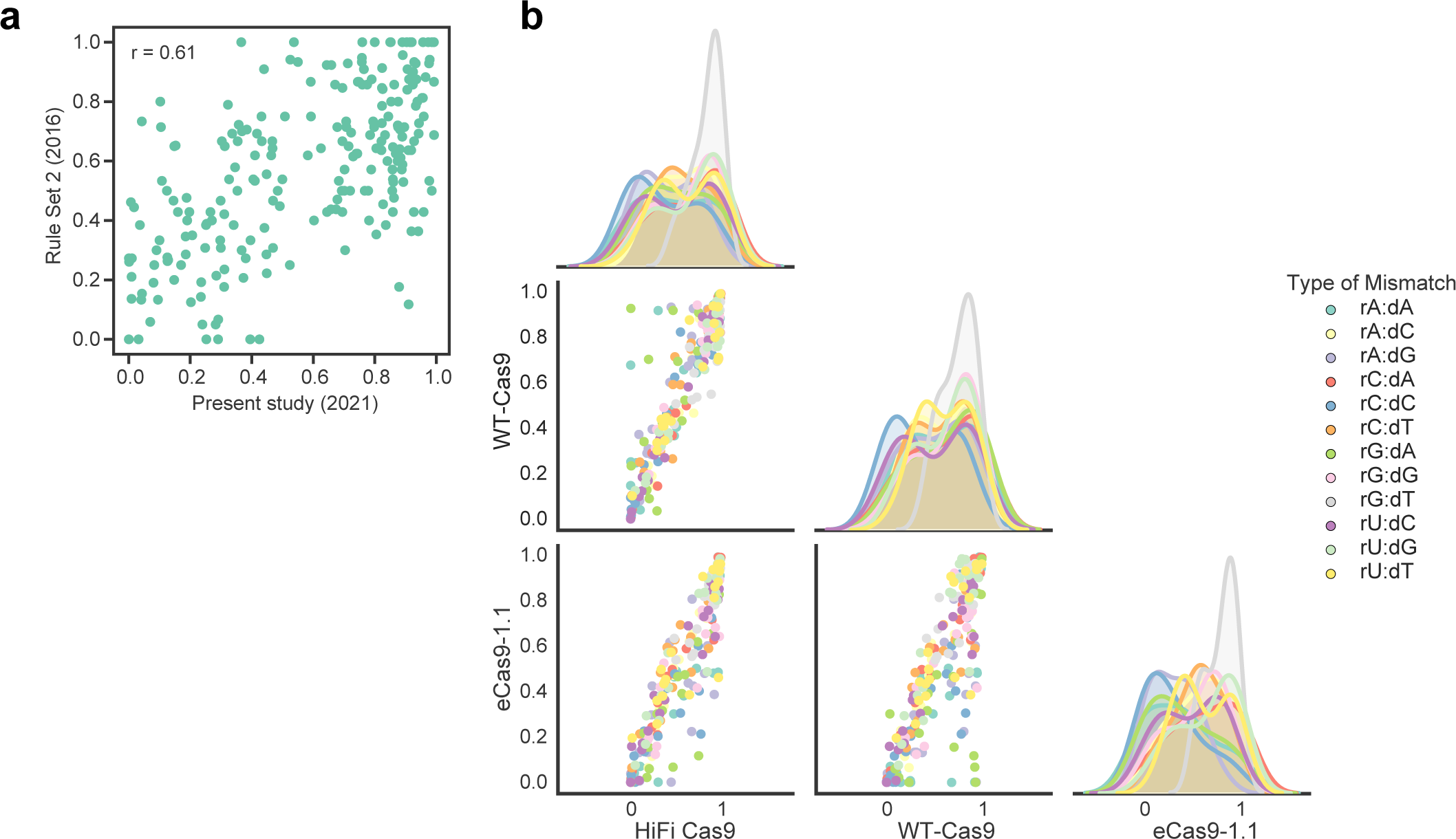
Off-target profiles of high fidelity variants. a) Comparison of CFD matrices for WT-Cas9 generated in this study (x-axis) and our previous work (y-axis). b) Comparison of the averaged probabilities of being active for each mismatch type between the Cas9 variants, shown as both scatter and kde plots.

**Supplementary Figure 3:**
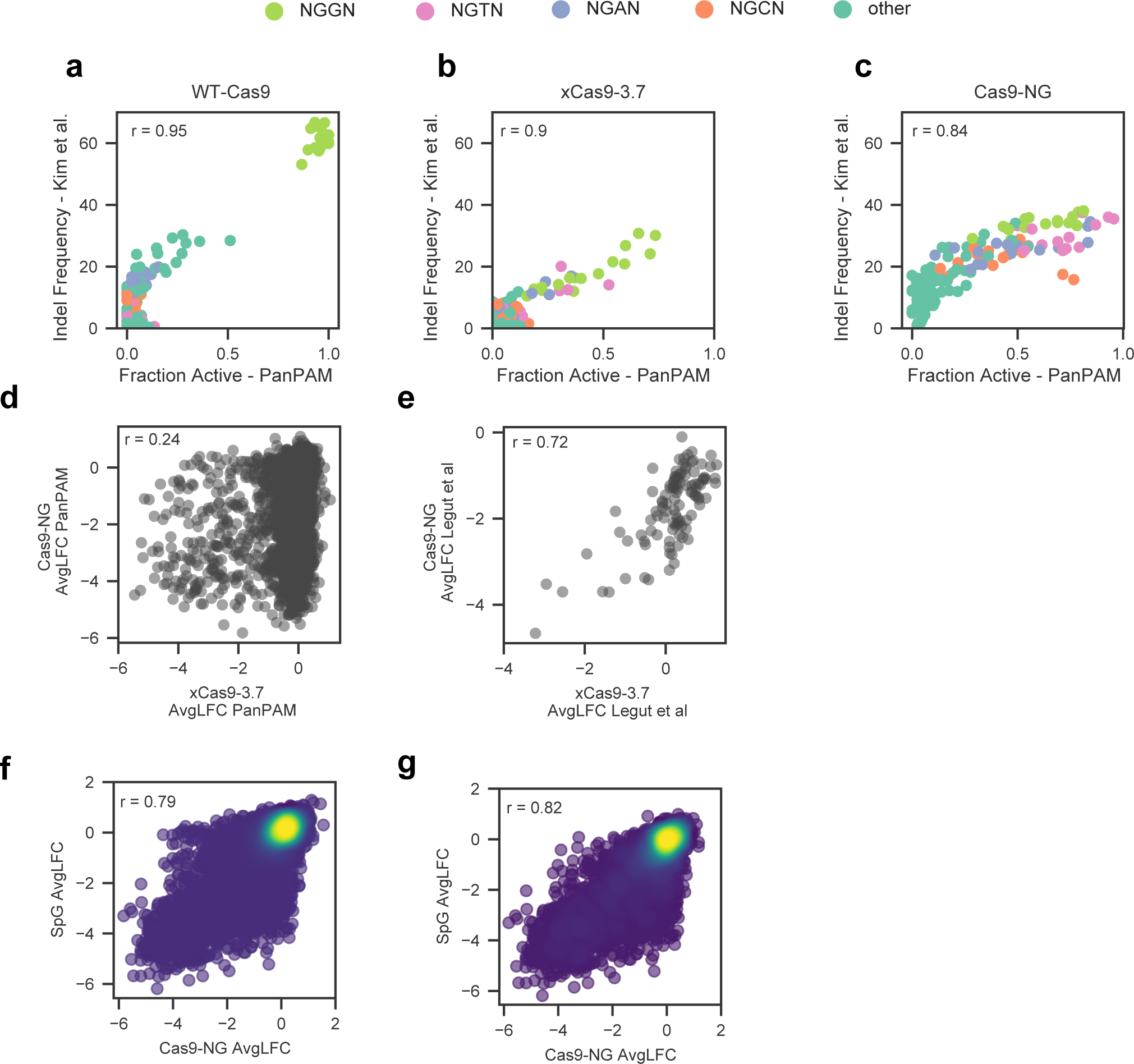
Benchmarking PAM-flexible variants. a,b,c) Comparison of fraction active values for WT-Cas9, xCas9-3.7 and Cas9-NG from the present PanPAM study (x-axis) and indel frequencies from Kim et al^31^ (y-axis). Each dot is a PAM. n = 148 PAMs. d) Comparison of avg LFC values for guides in the PAM-mapping library targeting essential genes screened with xCas9-3.7 and Cas9-NG. n = 2747 sgRNAs. e) Comparison of avg LFC values for guides targeting CD45 and CD55 described in Legut et al^24^. n = 108 sgRNAs. f) Comparison of sgRNAs screened with Cas9-NG and SpG. n = 18651 sgRNAs. g) Comparison of sgRNAs with an NG PAM screened with Cas9-NG and SpG. n = 4525 sgRNAs.

**Supplementary Figure 4:**
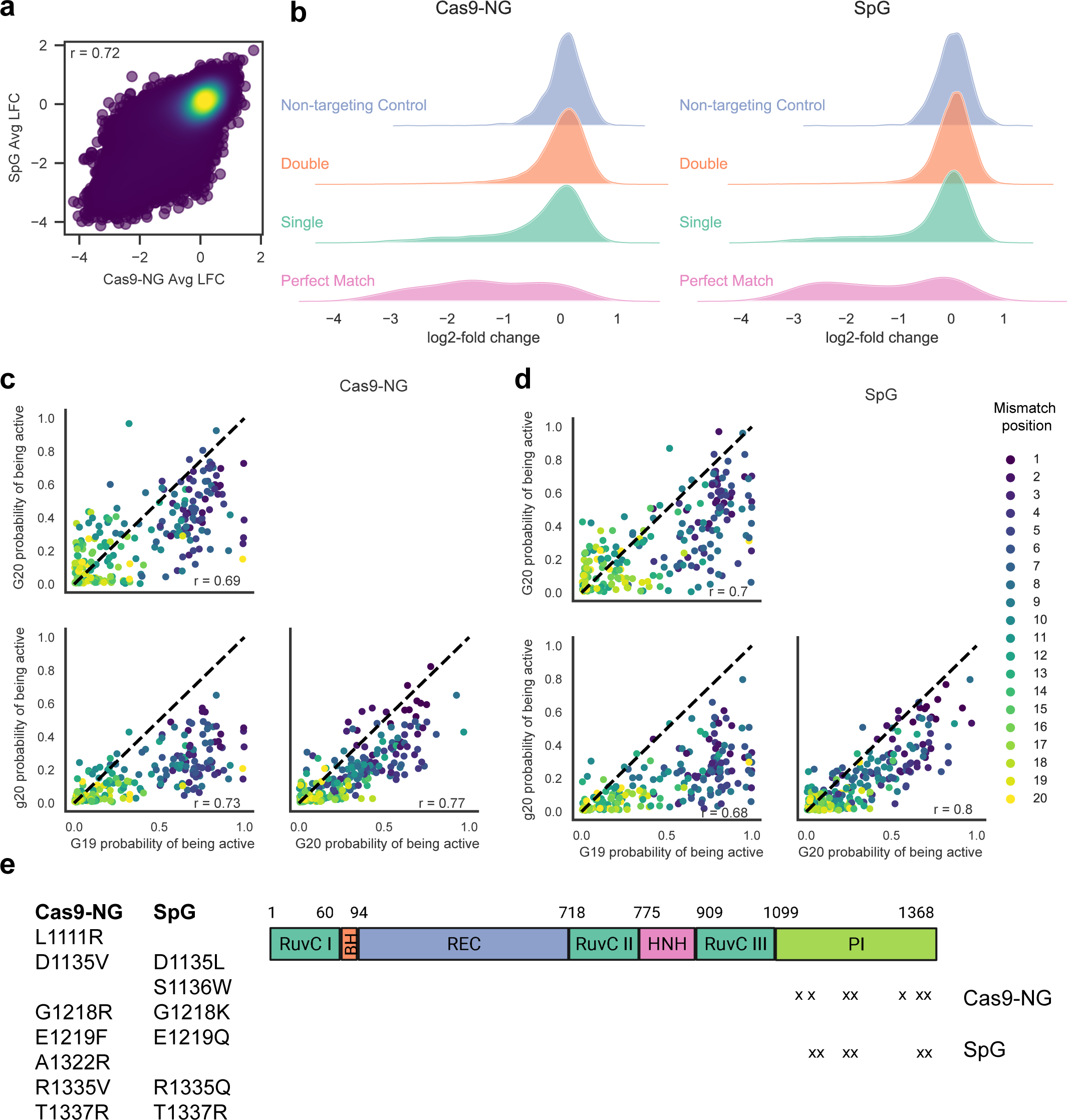
Off-target profiles of Cas9-NG and SpG. a) Correlation between Cas9-NG and SpG, screened with the off-target library. Pearson’s r is reported. n = 78058 sgRNAs. b) Ridge plots showing activity of unfiltered guides with zero, one or two mismatches. b) Scatter plots depicting the correlation between mismatches with sgRNAs of all 5’-types included in the library for Cas9-NG, colored by mismatch position. Each dot represents a type of mismatch at each position along the sgRNA (n = 228 pairings). d) Same as (c) but for SpG. e) Schematic depicting the mutations that result in Cas9-NG and SpG.

**Supplementary Figure 5:**
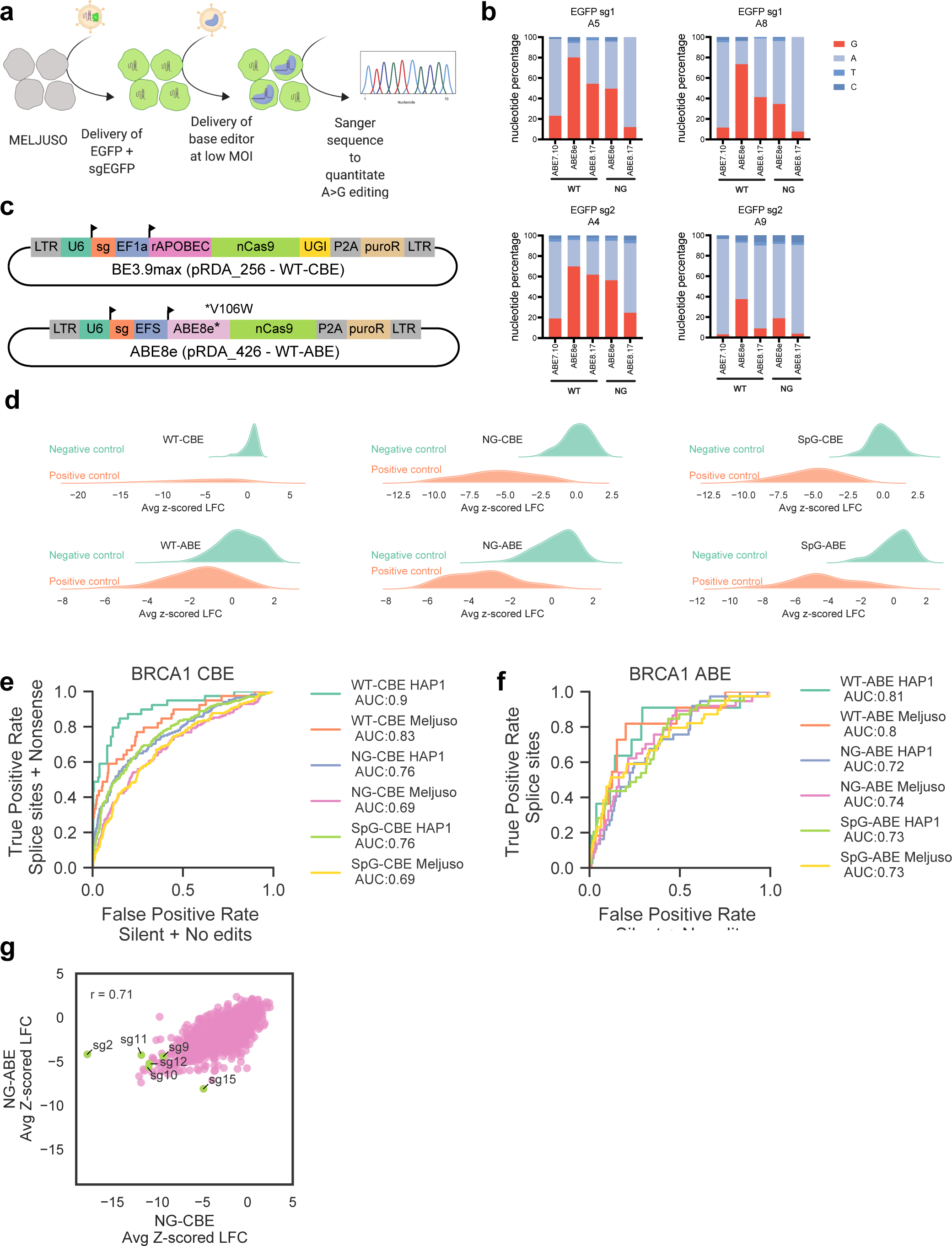
Tiling *BRCA1* with variant base editors. a) Schematic of EGFP activity assay. b) Nucleotide percentage at editable A’s in the sgRNA (denoted in the title of each barplot). c) Schematic of WT-CBE (top) and WT-ABE8e (bottom). Variant base editors were generated by introducing point mutations into the nCas9 of each vector. d) Ridge plots showing the distributions of positive and negative controls for the WT and variant base editors. e) ROC plot for each cell line screened with each Cas-CBE pairing. The AUC is reported. f) Same as (e) but for ABEs. Note that only splice sites are considered as true positives, as nonsense mutations cannot be introduced with ABE. g) Comparison of z-scored LFCs of guides predicted to make either a missense mutation with an NG-CBE and no change with an NG-ABE, no change with CBE and a missense mutation with ABE or a missense mutation with both base editors (n = 1694 sgRNAs). Guides that were selected for further validation are colored in green and labeled.

**Supplementary Figure 6:**
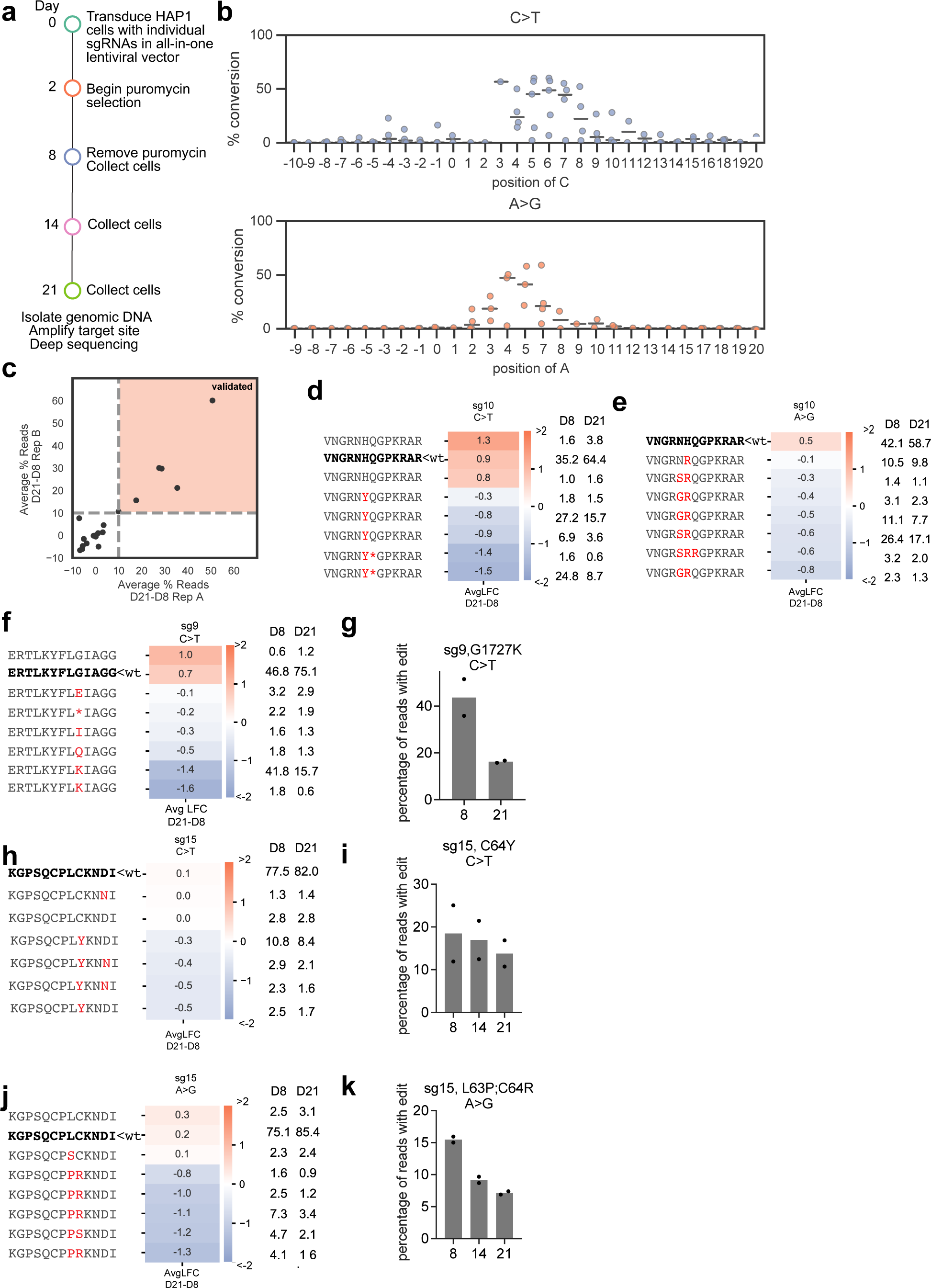
Validation of *BRCA1* hits identified by Cas9-NG. a) Timeline by which validation experiments were performed. b) C>T conversion (top) and A>G conversion (bottom) of guides with an intermediate/active PAM. Positions along the x-axis correspond with positions in the protospacer, where 1 corresponds to the first nucleotide of the protospacer and 21-23 correspond to the PAM. Lines show the median of all edits at that position across the validated sgRNAs. c) Validation scheme. sgRNAs in which the average % of reads increased by more than 10% from day 8 to day 21 are considered validated, as depicted with the orange shading. d,e,f,h,j) Translated sequence around the sgRNA for any allele with at least 1% abundance in any condition. The WT sequence is bolded in black, unchanged amino acids are in grey, and substitutions are highlighted in red. Avg LFC from day 21 - day 8 is indicated on the heatmap and relative percent abundance of each allele is indicated to the right (normalized after filtering for alleles with <1% abundance at both timepoints). g,i,k) Percentage of all sequencing reads containing the indicated mutation at each timepoint. Dots indicate n=2 biological replicates.

**Supplementary Figure 7:**
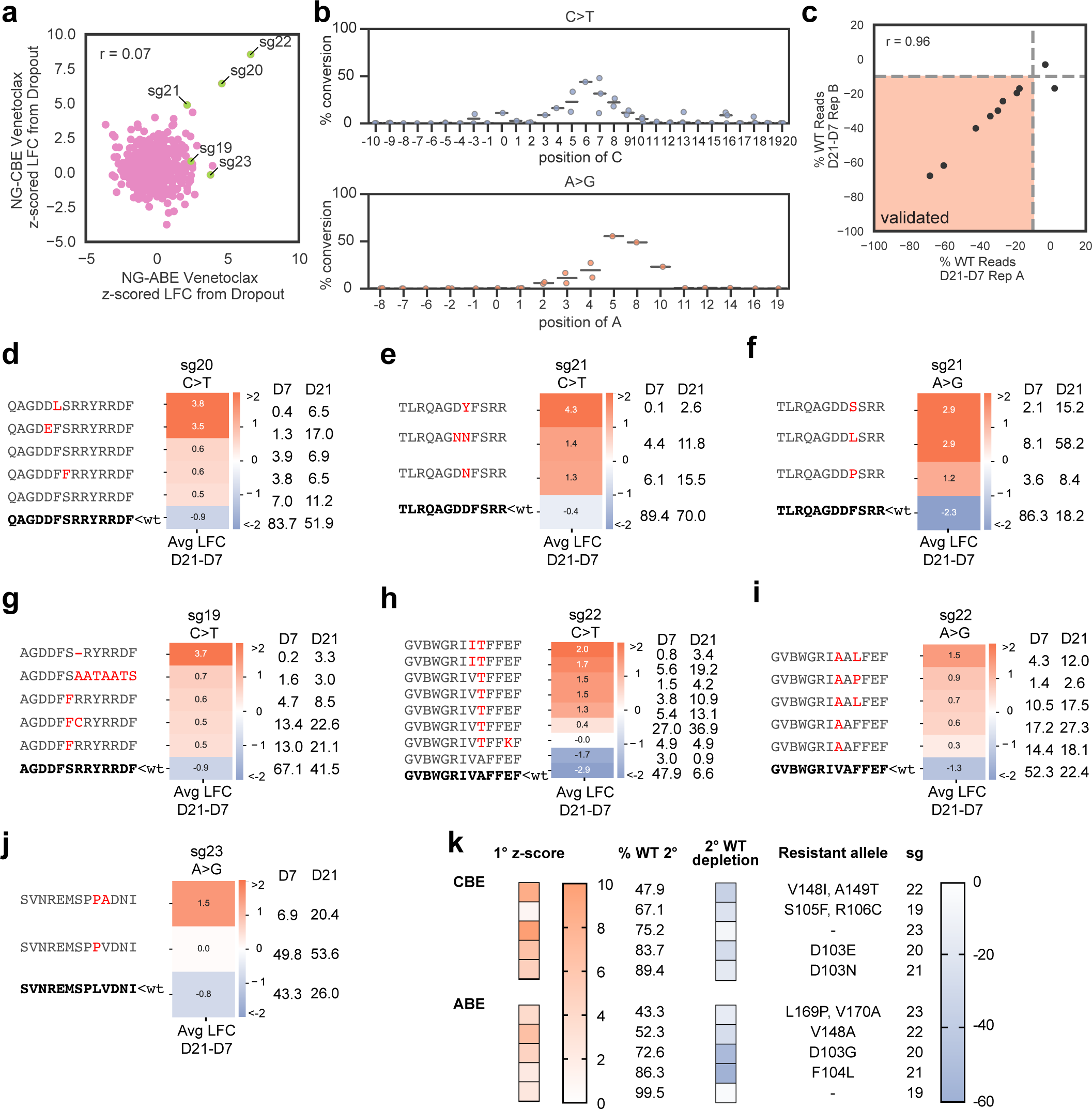
Validation of *BCL2* hits identified by Cas9-NG. a) Comparison of z-scored LFCs of guides predicted to make either a missense mutation with one base editor and a silent edit or no edit with the other, or a missense mutation with both base editors (n = 539 sgRNAs). Guides that were selected for further validation are colored in green and labeled. Pearson’s r is reported. b) C>T conversion (top) and A>G conversion (bottom) with sgs 19-23. Positions along the x-axis correspond with positions in the protospacer, where 1 corresponds to the first nucleotide of the protospacer and 21-23 correspond to the PAM. Lines show the median of all edits at that position across sgs 19-23. c) Validation scheme. sgRNAs in which the average % of WT reads depletes by more than 10% (under selective pressure of Venetoclax) from day 7 to day 21 are considered validated, as depicted with the orange shading. d-j) Translated sequence around the sgRNA for any allele with at least 1% abundance in any condition. The WT sequence is bolded in black, unchanged amino acids are in grey, and substitutions are highlighted in red. Avg LFC from day 21 - day 7 is indicated on the heatmap and relative percent abundance of each allele is indicated to the right (normalized after filtering for alleles with <1% abundance at both timepoints). k) Summary of validation results. 1° z-score indicates the average z-scored LFC of the sgRNA in the primary screen. % WT 2° indicates the % of reads that were still WT (unedited) on day 7 in the validation experiment. 2° WT depletion indicates the average change in the abundance of the WT allele from day 7 to day 21 in the Venetoclax-treated arm of the validation experiment.

## SUPPLEMENTAL DATA

Supplementary Data 1: PAM-mapping counts, library annotation, replicate correlations. Associated with **Figs 1,3.**

Supplementary Data 2: HF-off-target counts, library annotation, replicate correlations. Associated with Fig 2.

Supplementary Data 3: variant off-target counts, library annotation, replicate correlations. Associated with Fig 4.

Supplementary Data 4: LFC and probability of being active calculations for all of the off-target datasets that are used to calculate the CFD scores. Associated with **Figs 2,4**.

Supplementary Data 5: *BRCA1* CBE data - WT, NG, SpG counts, WT library annotation (includes different controls), variant library annotation, replicate correlations. Associated with Fig 5.

Supplementary Data 6: *BRCA1* ABE data - WT, NG, SpG counts, WT library annotation (includes different controls), variant library annotation, replicate correlations. Associated with Fig 5.

Supplementary Data 7: *BCL2* data - NG-CBE and NG-ABE counts, library annotations, replicate correlations. Associated with Fig 7.

Supplementary Data 8: Primers and guide sequences used for validation experiments and the parameters used to run all validation samples in CRISPResso2. Associated with **Figs 6,7**.

## Notes

https://www-ncbi-nlm-nih-gov.ezproxy.u-pec.fr/geo/query/acc.cgi?acc=GSE180351

